# Pharmacologic Targeting of ZNF281 Suppresses Metastatic Prostate Cancer Beyond Androgen Receptor Dependence

**DOI:** 10.64898/2026.09.03.749302

**Authors:** Guocheng Huang, Maria Areli Lorenzana-Carrillo, Hua Chen, Runtai Chen, Saba Abbasi Dezfouli, Joseph Nanoa, Dania Rimawi, Alois Haromy, Yuan-Yuan Zhao, Rohan Mittal, John Lewis, John R. Ussher, Evangelos D. Michelakis, Ronald B. Moore, Seyed Amirhossein Tabatabaei Dakhili, Gopinath Sutendra, Adam Kinnaird

## Abstract

Metastatic prostate cancer remains a lethal disease, with most treatments focusing on the androgen receptor (AR) axis. We identified the zinc finger protein 281 (ZNF281) as a previously unrecognized driver of metastatic prostate cancer that promotes both AR-related transcriptional output and distinct AR-independent tumor-promoting pathways. It increases AR expression and acts as a coactivator to promote AR transcriptional activity. Independent of AR, ZNF281 promotes prostate cancer by upregulating SMURF1 to sustain tumor growth and by promoting SNAIL-mediated metastasis. We developed an orally bioavailable, ZNF281 Interfering Molecule (Oral ZIM) that disrupts its DNA binding and AR protein interaction. Knockout of ZNF281 as well as treatment with oral ZIM potently inhibited prostate cancer growth and metastasis in orthotopic xenograft models of castration-sensitive and castration-resistant prostate cancers (without detectable systemic toxicity), and ZIM outperformed enzalutamide treatment in patient-derived prostate cancer organoids.

**Significance:** This study reveals the previously uncharacterized of ZNF281 in promoting androgen receptor–dependent transcription and androgen receptor-independent proliferative and metastatic programs in prostate cancer, advances understanding of disease progression across different androgen receptor expression statuses, and establishes pharmacologic ZNF281 inhibition as a therapeutic strategy for metastatic prostate cancer.

## INTRODUCTION

Metastatic prostate cancer remains an incurable disease. Five-year survival rates of prostate cancer decrease from 99% for localized disease to 31% in metastatic disease(1). Furthermore, there has been an approximate 5% annual increase in metastatic disease since 2011(2). Therefore, novel treatments are urgently needed. Inhibition of androgen receptor (AR) signaling (i.e., androgen deprivation therapy, ADT) remains the cornerstone of prostate cancer treatment. However, prolonged suppression of AR imposes a strong selective pressure that favors the emergence of more aggressive tumor cell populations capable of sustaining disease progression despite diminished or absent AR activity, ultimately contributing to castration-resistant prostate cancer (CRPC)(3–5). Nearly all patients eventually develop resistance to ADT, and treatment options specifically targeting metastatic lesions remain limited(6). Therefore, a therapeutic strategy that simultaneously targets AR-dependent signaling and AR-independent tumor-promoting pathways that remain active irrespective of AR status in metastatic prostate cancer may offer a survival benefit.

The Krüppel-type zinc-finger protein 281 (ZNF281) is a transcription factor implicated in regulating gene expression programs that facilitate epithelial-to-mesenchymal transition (EMT), cancer cell stemness, and purinergic signaling cascades(7,8). Its role in promoting cancer progression and metastasis has been demonstrated across multiple cancer types, including colorectal cancer, hepatocellular carcinoma, pancreatic cancer, melanoma, lung cancer and breast cancer(7,9–13). However, the potential functions of ZNF281 in metastatic prostate cancer remain unclear. Intriguingly, unbiased mass spectrometry has identified ZNF281 as an AR-interacting protein, suggesting it may have a role in AR-mediated signaling in prostate cancer(14). Here, we show that ZNF281 is a coactivator for the AR, enhancing its transcriptional activity, while also regulating its expression. We found that ZNF281 has an important role in the progression of PCa in an AR-dependent and AR-independent manner. To provide a therapeutic option, we synthesized a novel small-molecule inhibitor named ZNF281 Interfering Molecule (ZIM). The mechanism of action and selectivity of the first version ZIM have been previously characterized(13). In the present study, we developed a structurally optimized ZIM containing fewer hydrophobic groups. This optimized ZIM supported oral dosing in preclinical *in vivo* studies while retaining its biological activity against ZNF281. Oral ZIM treatment disrupts ZNF281-associated transcriptional activities and attenuates its AR-coactivator functions, leading to significant anti-tumor effects in orthotopic models of metastatic castration-sensitive and castration-resistant prostate cancer, as well as in patient-derived prostate cancer organoids.

## RESULTS

### ZNF281 is expressed in metastatic prostate cancer, and its inhibition suppresses tumor growth and metastasis

We examined the protein levels of ZNF281 in PCa. Paired tissue samples from 15 patients with metastatic prostate cancer who underwent radical prostatectomy with bilateral lymph-node dissection (**Supplementary Table S1**) were subjected to immunofluorescence staining to assess ZNF281 expression in lymph node metastases, primary tumors, and adjacent normal prostate tissues (**Fig. 1A**). Quantification revealed that metastatic lesions exhibited significantly higher ZNF281 protein levels compared with primary tumors and normal tissues (**Fig. 1B**). Stratification by Gleason Grade Group (GGG) demonstrated that higher-grade tumors (GGG 3-5) expressed significantly greater amounts of ZNF281 than lower-grade tumors (GGG 1-2) (**Fig. 1C**). Moreover, ZNF281 expression in primary tumors positively correlated with its levels in matched metastatic tissues (r = 0.63, *P* = 0.0141, **Fig. 1D**). Analysis of 489 prostate adenocarcinoma patients’ survival data from TCGA database showed that high *ZNF281* copy number alteration (CNA) is significantly related to poor progression-free survival of the patients (**Fig. 1E**). Together, these data indicate that ZNF281 is induced during prostate cancer progression.

**Figure 1.**
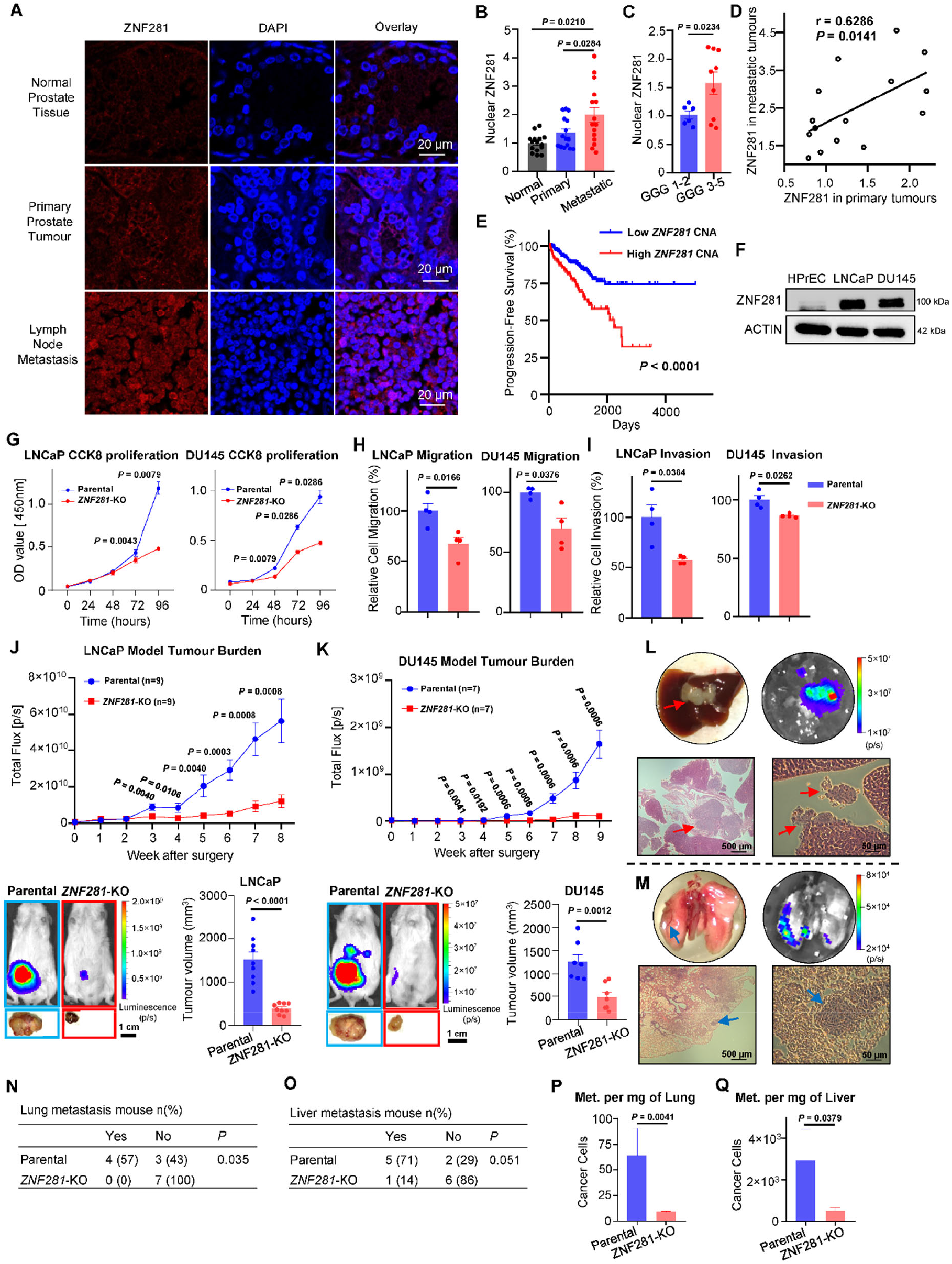
ZNF281 is expressed in metastatic prostate cancer, and its inhibition suppresses tumor growth and metastasis. **A,** Representative immunofluorescence images of ZNF281 in paired tissue sections from prostate cancer patients. **B,** Quantitative analysis of 15 patients showing nuclear ZNF281 expression in metastatic tissues, primary tumors and normal prostate tissues. **C,** Stratified analysis revealing ZNF281 expression in low-grade tumors (Gleason Grade Group 1-2) and high-grade prostate cancers (Gleason Grade Group 3-5). **D,** ZNF281 protein expression levels in normal prostate epithelial cells (HPrEC) versus metastatic prostate cancer cell lines (LNCaP and DU145). **E,** Correlation analysis indicating the relationship between ZNF281 expression in primary tumors and its expression in matched metastatic lesions. **F,** Kaplan-Meier survival analysis between the ZNF281 copy number alteration (CNA) and progression-free survival in 489 prostate adenocarcinoma patients (figure modified from https://www.tcga-survival.com/). **G,** *In vitro* CCK-8 cell proliferation assay in both LNCaP and DU145 cells. **H,** Transwell migration assay in LNCaP and DU14. **I,** Transwell invasion assay shows that ZNF281-KO LNCaP and DU145 cells exhibited reduced invasive capacity. **J,** (LNCaP)**, K,** (DU145) Statistical data and representative images of the orthotopic xenograft mouse models. Tumor growth *in vivo* was monitored weekly by bioluminescence imaging. ZNF281-KO tumors grew significantly more slowly and exhibited smaller endpoint tumor volumes, and showed fewer metastatic lesions in DU145 model. **L, M,** Representative images of the live and lung metastatic lesions in the DU145 orthotopic xenograft mouse model. Liver metastases (red arrows) and lung metastases (blue arrows) were further confirmed with tumor cell-specific luminescence signals and H&E staining. **N, O,** Statistical analysis of mice with lung metastasis and mice with liver metastasis. **P, Q,** Quantification of tumor burden in lung metastases and in liver metastases. Graphs show the mean ± SEM from three independent biological replicates, or number (%). Statistical significance was determined using RM one-way ANOVA with Tukey’s post hoc multiple-comparison test **(B),** *t* test with Welch’s correction **(C, G, H, I)**, Spearman’s correlation analysis (**D**), Kaplan–Meier survival analysis (**E**), Mann–Whitney test **(J, K, P, Q)**, or one-sided Fisher’s exact test **(N, O)**.

To explore the functional significance of ZNF281 in metastatic prostate cancer, we utilized two metastatic PCa cell lines, LNCaP (AR-positive) and DU145 (AR-negative), both of which exhibited markedly elevated ZNF281 protein levels relative to normal prostate epithelial cells (**Fig. 1F**). Efficient depletion of ZNF281 was observed in both ZNF281-knockout (KO) cell lines by immunofluorescence staining (**Supplementary Fig. S1A, B**). ZNF281 KO in LNCaP and DU145 cells impaired cell proliferation (**Fig. 1G**), as well as transwell-based migration (**Fig. 1H**) and invasion (**Fig. 1I**), compared to parental cells. These data suggest that ZNF281 may have an important role for prostate cancer proliferation and motility.

To provide translational relevance to our approach, we generated an orthotopic xenograft mouse model for human prostate cancer. Consistent with the *in vitro* findings, immunofluorescence staining confirmed persistent loss of ZNF281 expression in orthotopic tumors derived from ZNF281-KO LNCaP and DU145 cells (**Supplementary Fig. S1C, D**). We compared parental to ZNF281-KO LNCaP or DU145 tumors and found a robust decrease in tumor growth in the ZNF281-KO groups (**Fig. 1J, K**). In the DU145 cohort, prominent metastatic lesions were observed in the liver (**Fig. 1L**) and lungs of parental mice (**Fig. 1M**). ZNF281 knockout eliminated lung metastasis (57% vs. 0%, *P* = 0.035, **Fig. 1N**), and showed a trend toward reducing the incidence of liver metastasis (71% vs. 14%, *P* = 0.051, **Fig. 1O**), although the latter difference didn’t reach statistical significance compared with the parental group. It also decreased the tumor burden at the metastatic sites (**Fig. 1P, Q**).

These results demonstrate that ZNF281 promotes tumor growth and metastasis in both AR-positive (LNCaP) and AR-negative (DU145) models, suggesting that ZNF281 may regulate prostate cancer progression regardless of the AR status. Given this possibility, we sought to elucidate the molecular basis of ZNF281 function.

### ZNF281 regulates AR expression in prostate cancer by increasing AR transcription

To investigate the potential mechanism of ZNF281 in prostate cancer, we analyzed a cohort of 493 primary prostate adenocarcinoma samples from cBioPortal. Patients were stratified based on ZNF281 mRNA expression (z-score): the top 65 patients (≥ 1 SD) were defined as the high-ZNF281 group, whereas the bottom 64 patients were defined as the low-ZNF281 group. Differentially expressed genes (DEGs) analysis revealed AR as the second most increased gene among 1,760 significantly upregulated transcripts (fold change = 10.2) (**Fig. 2A**). Pearson correlation analysis further demonstrated a strong positive correlation between ZNF281 and AR mRNA expression (r = 0.73, *P* < 0.0001, **Fig. 2B**). Together, these data indicate a close association between ZNF281 and AR expression.

**Figure 2.**
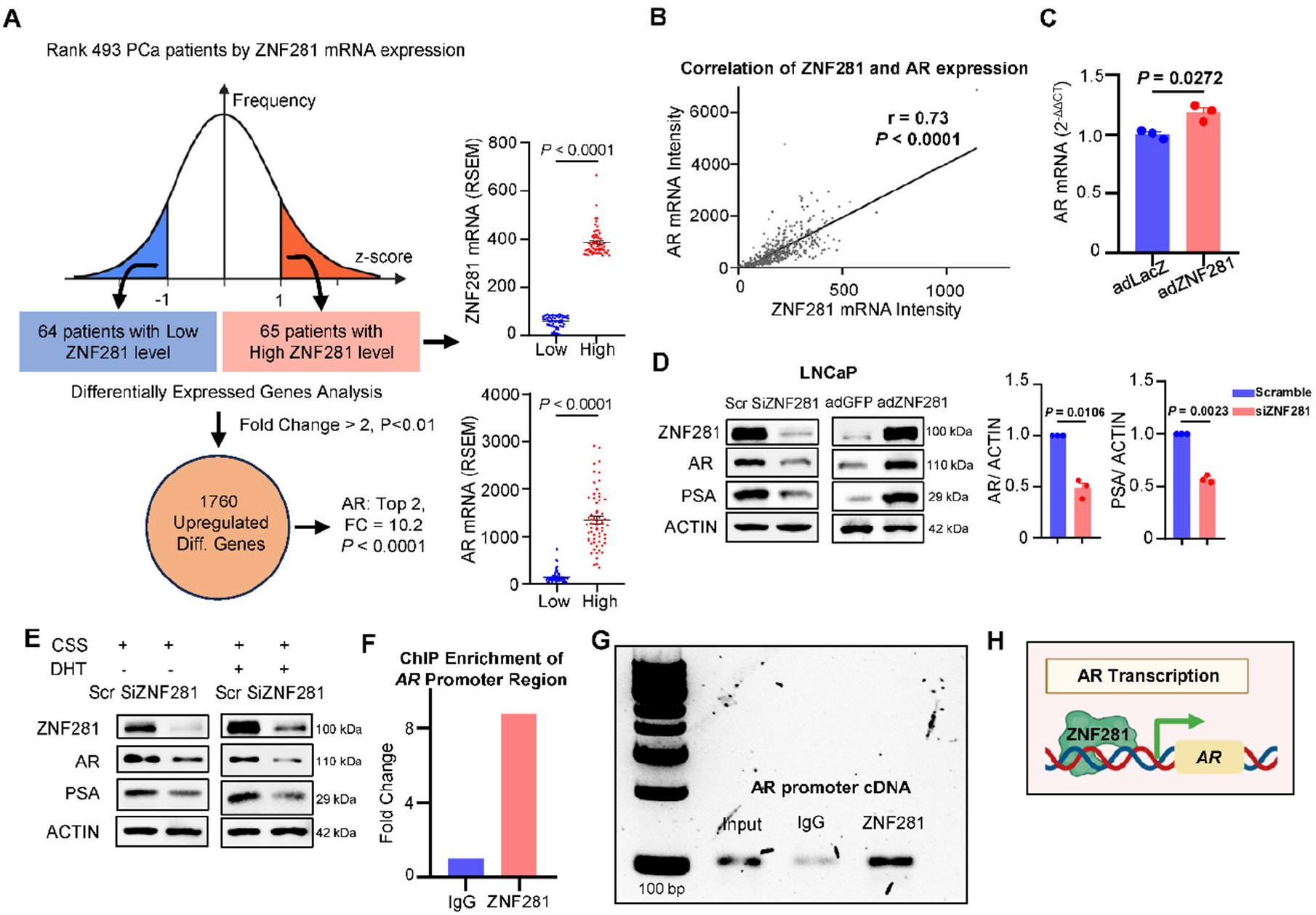
ZNF281 regulates AR expression in prostate cancer by increasing AR transcription. **A,** Differentially expressed genes analysis between high and low ZNF281 expression groups in a cohort of 493 primary prostate adenocarcinoma samples (cBioPortal). **B,** Correlation analysis between ZNF281 and AR mRNA expression levels. **C,** AR mRNA expression in LNCaP cells following ZNF281 overexpression. **D,** Western blot analysis and quantification showing the effects of ZNF281 manipulation on the protein levels of AR and its target gene PSA. **E,** Western blot analysis shows that ZNF281 knockdown reduced the protein expression of AR and its target gene PSA, with or without dihydrotestosterone (DHT) treatment. **F,** ChIP–qPCR analysis showing enrichment of ZNF281 at the AR promoter region. **G,** Agarose gel electrophoresis showing a single ChIP–qPCR product of the expected size. **H,** Schematic illustration of the regulatory role of ZNF281 in AR expression. Graphs show the mean ± SEM from three independent biological replicates. Statistical significance was determined using the *t* test **(A)**, *t* test with Welch’s correction **(C, D)**, and Pearson’s correlation analysis **(B)**.

Since both ZNF281 and AR function as transcription factors, we next explored their possible regulatory relationship. ZNF281 expression was depleted using siRNA or overexpressed using adenoviral vectors in LNCaP cells. ZNF281 overexpression significantly increased AR mRNA levels compared to control-infected LNCaP cells (**Fig. 2C**). Consistently, we found that ZNF281 knockdown in LNCaP cells markedly reduced AR levels, along with its downstream target PSA, whereas ZNF281 overexpression had the opposite effect compared to knockdown **(Fig. 2D**). In contrast, AR knockdown did not decrease ZNF281 levels (**Supplementary Fig. S2A**). A similar regulatory effect of ZNF281 overexpression on AR mRNA (**Supplementary Fig. S2B**) and protein levels (**Supplementary Fig. S2C**) was also observed in the immortalized non-cancer prostate epithelial cell line RWPE1.

Moreover, inhibition of ZNF281 decreased the protein levels of AR and its target gene PSA, under both steroid-depleted (charcoal-stripped FBS) and dihydrotestosterone (DHT)-stimulated conditions, indicating that the regulatory effect of ZNF281 on AR was observed irrespective of androgen availability (**Fig. 2E**). ZNF281 ChIP-qPCR (**Fig. 2F**) demonstrated significant enrichment of ZNF281 at the AR promoter region, and agarose gel electrophoresis revealed a single qPCR product of the expected size (**Fig. 2G**), supporting the specificity of the ChIP-qPCR product. These findings support a role for ZNF281 in regulating AR transcription (**Fig. 2H**).

### ZNF281 is a coactivator for AR

We next investigated whether ZNF281 also regulates AR transcriptional activity through association with the AR protein. Co-immunoprecipitation of endogenous ZNF281 in LNCaP cell lysates detected AR protein in ZNF281 immunoprecipitates (**Fig. 3A**). These findings suggested that ZNF281 may associate with the AR in protein level.

**Figure 3.**
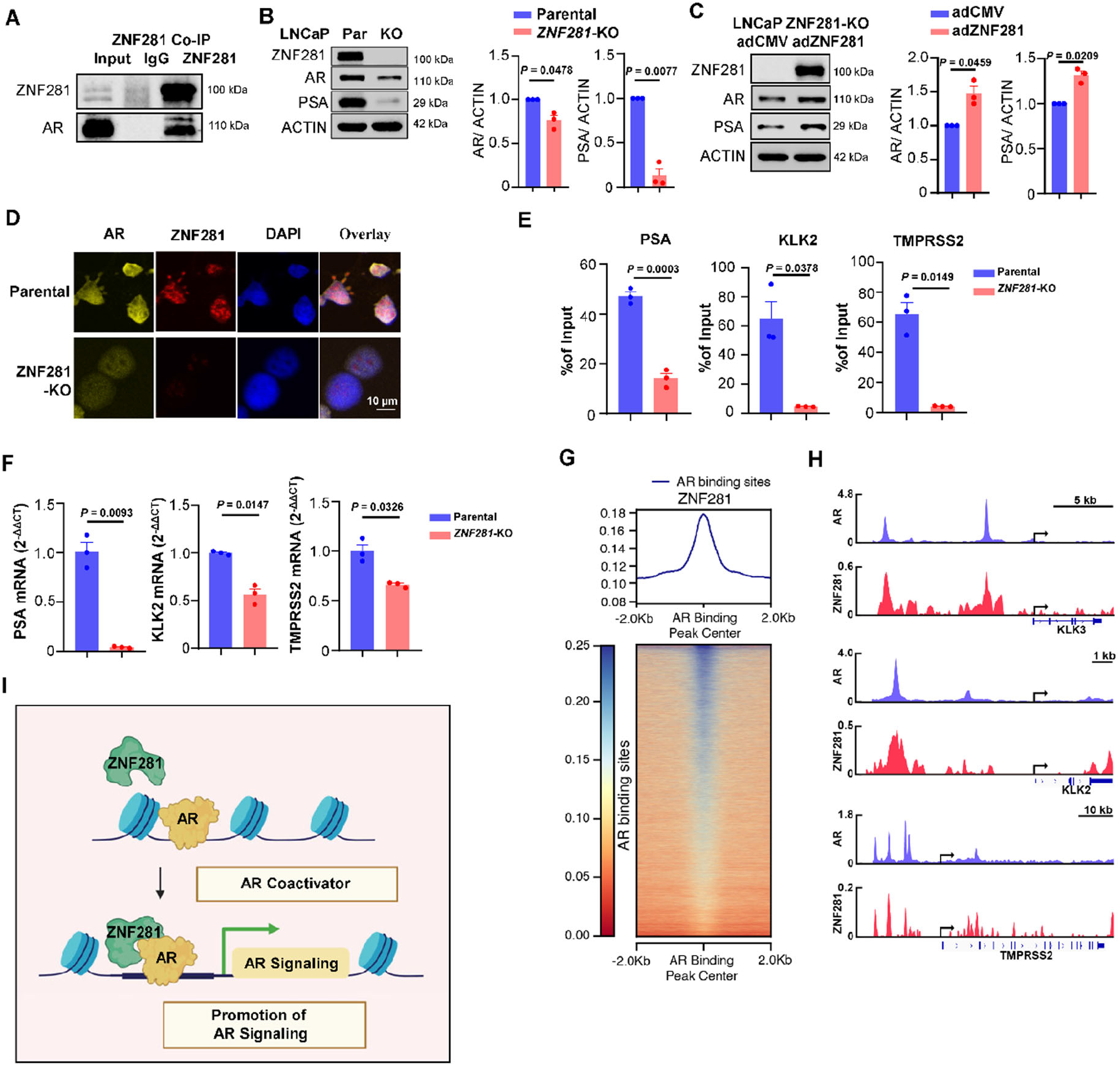
ZNF281 is a coactivator for AR. **A,** Co-immunoprecipitation analysis in LNCaP cells showing that ZNF281 may associate with AR protein. **B,** Western blot analysis and quantification showing the effects of ZNF281 knockout (KO) in LNCaP cells. **C,** Western blot analysis and quantification showing the effects of ZNF281 re-expression in ZNF281-KO LNCaP cells. **D,** Representative immunofluorescence images showing AR expression and intracellular localization in ZNF281-KO LNCaP cells. **E,** ChIP–qPCR analysis of AR in parental and ZNF281-KO LNCaP cells showing reduced AR binding to its target gene loci in ZNF281-KO cells, including PSA, KLK2, and TMPRSS2. **F,** RT–qPCR analysis showing mRNA expression levels of AR target genes in parental and ZNF281-KO LNCaP cells. **G,** Metaprofile (top) and heatmap (bottom) showing ZNF281 ChIP-seq signal centered on AR binding sites (±2 kb) identified by AR ChIP-seq in LNCaP cells. The average ZNF281 signal is enriched at the center of AR peaks, indicating preferential localization of ZNF281 at AR-occupied regulatory regions. **H,** Genome browser tracks showing representative examples of AR (blue) and ZNF281 (red) ChIP-seq signal at canonical AR target loci, including KLK3, KLK2, and TMPRSS2. Gene models are shown below, with arrows indicating transcriptional orientation. Scale bars indicate genomic distances as noted. **I,** Schematic illustration of the coactivator role of ZNF281 in AR transcriptional activity. Graphs show the mean ± SEM from three independent biological replicates. Statistical significance was determined using *t* test with Welch’s correction **(B, C, E, F)**.

Consistent with this, ZNF281-KO LNCaP cells exhibited a modest decrease in AR protein levels but a marked reduction in PSA expression compared to parental cells (**Fig. 3B**). Re-expression of ZNF281 in the KO cells partially restored the levels of AR and PSA (**Fig. 3C**). Although ZNF281 knockout did not alter AR nuclear localization (**Fig. 3D**), ChIP-qPCR revealed a significant reduction in AR occupancy at the promoter regions of its target genes, including PSA (KLK3), TMPRSS2, and KLK2 (**Fig. 3E**), accompanied by a corresponding decrease in their mRNA levels (**Fig. 3F**). ChIP-seq analysis further confirmed the overlapping occupancy of ZNF281 and AR at regulatory regions (**Fig. 3G**), including canonical AR target loci PSA, KLK2, and TMPRSS2 (**Fig. 3H**).

Collectively, these findings demonstrate that ZNF281 depletion not only inhibits AR expression but also impairs its transcriptional activity. ZNF281 regulates AR’s expression and may function as a coactivator required for optimal AR-mediated transcription in prostate cancer cells (**Fig. 3I**).

### AR-independent functions of ZNF281 in prostate cancer progression involve SMURF1-mediated growth and SNAIL-mediated EMT

To investigate the AR-independent mechanism of ZNF281, we performed unbiased RNA-seq and ChIP-seq in DU145 (AR-negative) and LNCaP (AR-positive) cells. A total of 94 overlapping DEGs were identified between the two cell lines (**Fig. 4A**). Apart from the robust repression of the androgen receptor pathway in the LNCaP ZNF281-KO cells compared to parental cells, PROGENy pathway analysis of both LNCaP and DU145 revealed activation of the p53 pathway and repression of the PI3K pathway upon loss of ZNF281 (**Fig. 4B**). Among the 94 shared DEGs, 17 were downregulated in ZNF281-KO LNCaP and ZNF281-KO DU145 cells (**Fig. 4C**). Integration analysis of ChIP-seq data showed that three of these genes (SMURF1, FARP2, and TEX2) were directly bound by ZNF281 in their promoter regions (**Fig. 4D, Supplementary Fig. S3A**).

**Figure 4.**
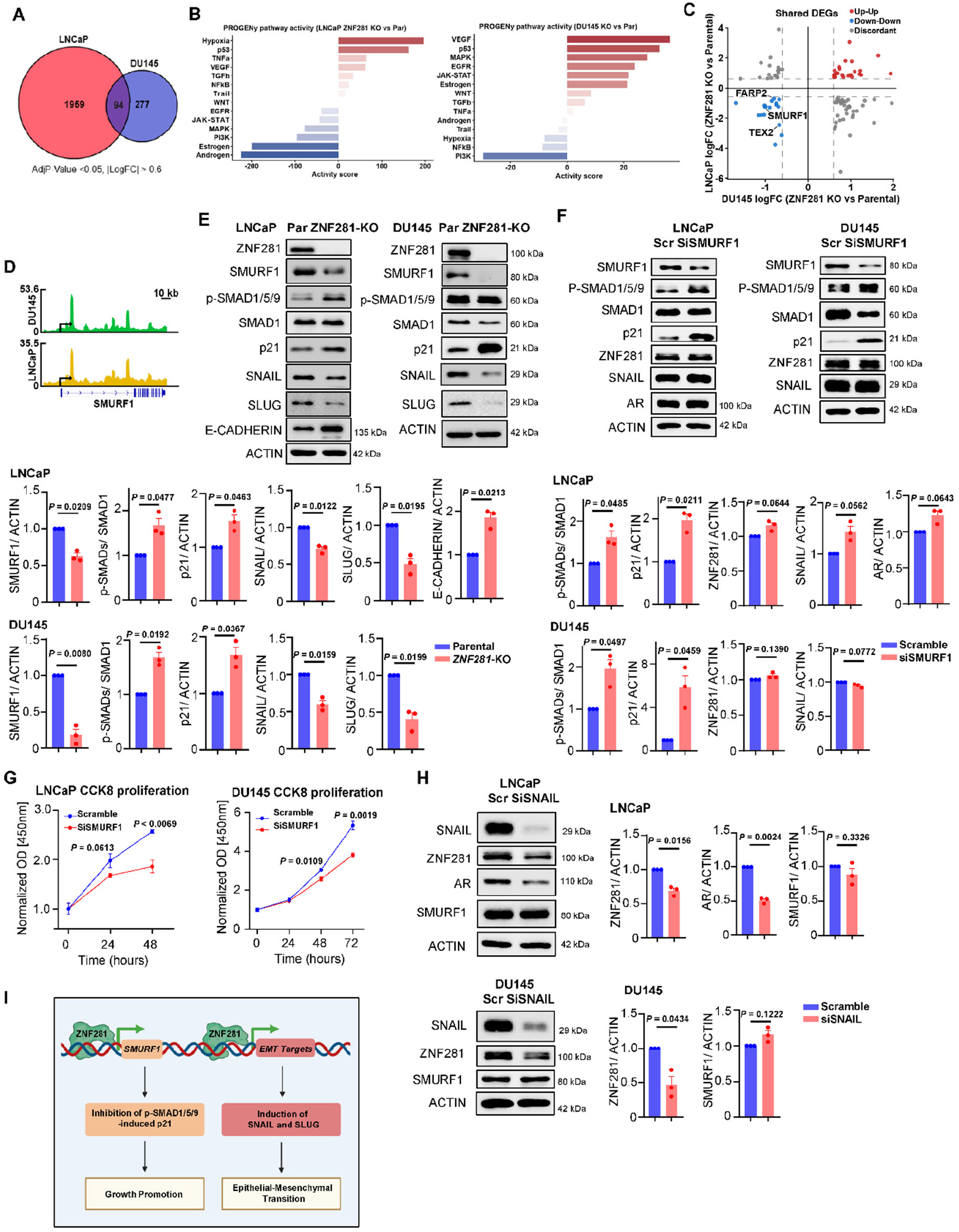
AR-independent roles of ZNF281 in prostate cancer progression involve SMURF1-mediated growth and SNAIL-mediated EMT. **A,** Venn diagram showing the overlap of differentially expressed genes (DEGs) following ZNF281 knockout in LNCaP and DU145 cells. DEGs were defined using an adjusted P value < 0.05 and |logFC| > 0.6. While the majority of transcriptional changes are cell-line specific, a subset of genes is commonly regulated in both contexts. **B,** PROGENy pathway activity analysis inferred from transcriptional changes in LNCaP and DU145 ZNF281-KO cells relative to their parental controls. Bar plot shows inferred activation (positive scores) or repression (negative scores) of major signalling pathways, revealing broad alterations in oncogenic and stress-associated signalling networks following ZNF281 loss. **C,** Scatter plot comparing log fold-changes (logFC) of shared DEGs between LNCaP (y-axis) and DU145 (x-axis) following ZNF281 loss. Genes are color-coded based on concordant regulation (upregulated in both cell lines, red; downregulated in both cell lines, blue) or discordant regulation (gray). Selected representative genes are highlighted. **D,** Genome browser tracks showing ZNF281 ChIP-seq signal in DU145 (green) and LNCaP (yellow) cells at the representative shared DEG SMURF1. Tracks illustrate differential ZNF281 promoter occupancy across cell lines. Gene models and transcriptional orientation are shown below, with scale bars indicating genomic distance. **E,** Western blot analysis and quantification showing the effects of ZNF281 knockout on SMURF1 signalling and EMT-related markers in LNCaP and DU145 cells. **F,** Western blot analysis and quantification showing the effects of SMURF1 inhibition by siRNA transfection on p-SMAD1/5/9–p21 axis and the expression of ZNF281, SNAIL and AR in LNCaP cells, and ZNF281 and SNAIL in DU145 cells. **G,** CCK-8 assays showing the effects of siRNA-mediated SMURF1 knockdown on the proliferation of LNCaP and DU145 cells. **H,** Western blot analysis and quantification showing the effects of siRNA-mediated SNAIL knockdown on ZNF281, AR and SMURF1 expression in LNCaP cells and on ZNF281 and SMURF1 expression in DU145 cells. **I,** Schematic illustration of the AR-independent mechanisms of ZNF281 in prostate cancer. Graphs show the mean ± SEM from three independent biological replicates. Statistical significance was determined using t test with Welch’s correction.

Among these three genes, SMURF1 (Smad ubiquitin regulatory factor 1) exhibited the highest absolute log fold change. It has been reported to promote cancer survival and progression through multiple signal pathways(15), including promoting p53 degradation and activating the PI3K/Akt/mTORC1 pathway(16,17). Moreover, SMURF1 is a negative regulator of TGFβ/BMP signaling by promoting p-SMAD1/5/9 degradation and is known to sustain cancer cell stemness(18). In contrast, activation of p-SMAD1/5/9 suppresses cancer cells’ proliferation by inducing the expression of p21(19,20). Specifically, functional enrichment analysis of DEGs in LNCaP highlighted mitotic cell cycle transition, whereas it highlighted epithelial cell proliferation in DU145 (**Supplementary Fig. S3B-E**).

We speculated that SMURF1 may be an important mediator for ZNF281-mediated AR-independent prostate cancer progression. Our sequencing data showed a significant reduction in SMURF1 mRNA expression in ZNF281-KO LNCaP and DU145 cells, which was further confirmed at the protein level (**Fig. 4E**). Similarly, siRNA-mediated knockdown of ZNF281 decreased SMURF1 protein expression, whereas re-expression of ZNF281 in the ZNF281-KO LNCaP cells rescued SMURF1 levels (**Supplementary Fig. S4A, B**). These data revealed the transcriptionally regulated effects of ZNF281 on SMURF1 in prostate cancer.

To determine the functional significance of SMURF1, we silenced SMURF1 using siRNA in both DU145 and LNCaP cells. SMURF1 knockdown increased the level of p-SMAD1/5/9 and p21(**Fig. 4F**), accompanied by significantly reduced CCK-8 proliferation (**Fig. 4G)**, indicating that SMURF1 promotes prostate cancer cell proliferation. Consistent with these findings, ZNF281-KO markedly increased p-SMAD1/5/9 relative to the total SMAD1 and elevated p21 compared to parental cells, indicating restoration of the p-SMAD1/5/9 mediated antiproliferative signaling in the ZNF281-KO cells (**Fig. 4E**). Conversely, re-expression of SMURF1 in ZNF281-KO cells suppressed p-SMAD1/5/9 signaling and restored proliferation in both ZNF281-KO LNCaP and DU145 cells (**Supplementary Fig. S4C, D**), supporting SMURF1 as a downstream mediator of the proliferative function of ZNF281.

ZNF281 has been reported to promote cancer metastasis by activating the transcription of *SNAIL*(*21,22*). Consistent with this, both ZNF281 knockout and siRNA-mediated knockdown of ZNF281 markedly reduced the EMT markers SNAIL and SLUG (**Fig. 4E, Supplementary Fig. S4A**). These effects were reversed by ZNF281 re-expression in LNCaP ZNF281-KO cells, which also suppressed E-cadherin expression (**Supplementary Fig. S4B**), further supporting the role of ZNF281 in promoting EMT.

Although both SMURF1 and SNAIL were regulated by ZNF281, it remained unclear whether they represent a common signaling cascade or distinct downstream pathways. We therefore investigated the relationship between the SMURF1 and SNAIL axes. Interestingly, siRNA-mediated SMURF1 knockdown did not alter AR expression in LNCaP cells or ZNF281 and SNAIL expression in either LNCaP or DU145 cells (**Fig. 4F**). Consistently, SMURF1 inhibition did not significantly affect the migration or invasion of either cell line (**Supplementary Fig. S4E, F**). In contrast, knockdown of SNAIL reduced ZNF281 expression and significantly impaired Transwell invasion in both LNCaP and DU145 cells (**Fig. 4H, Supplementary Fig. S4G**). These findings are consistent with previous evidence that SNAIL induces ZNF281 expression, whereas ZNF281 directly activates SNAIL, thereby establishing a reciprocal feed-forward regulatory circuit that promotes EMT and tumor progression(21). In LNCaP cells, SNAIL knockdown also reduced AR expression and impaired CCK-8 proliferation (**Fig. 4H, Supplementary Fig. S4H**). However, despite the reduction in ZNF281 expression, SNAIL knockdown did not significantly alter SMURF1 expression in either LNCaP or DU145 cells, nor did it affect CCK-8 proliferation in DU145 cells (**Fig. 4H, Supplementary Fig. S4I**). This suggests that the reduction in ZNF281 induced by SNAIL knockdown may be insufficient to suppress the ZNF281-SMURF1 regulatory axis. Moreover, the differential proliferative effects observed in AR-positive LNCaP and AR-negative DU145 cells indicate that the contribution of SNAIL to prostate cancer cell proliferation maybe context-dependent.

The regulation of both SMURF1 and SNAIL by ZNF281 was further validated in another AR-negative metastatic prostate cancer cell line, PC3 (**Supplementary Fig. S4J**). Collectively, these findings indicate that ZNF281 may regulate two functionally distinct downstream programs: the SMURF1–p-SMAD1/5/9 axis predominantly promotes prostate cancer cell proliferation, whereas the SNAIL pathway primarily drives EMT and invasion. The lack of reciprocal regulation between SMURF1 and SNAIL suggests that these two pathways may operate largely in parallel downstream of ZNF281, highlighting ZNF281 as a potential regulator of both proliferative and metastatic programs in prostate cancer.

In summary, ZNF281 may promote prostate cancer progression independently of AR by <u>1.</u> Directly upregulating SMURF1, thereby suppressing the antiproliferative effect of p-SMAD1/5/9; <u>2.</u> Activating EMT-promoting transcription, including SNAIL expression. This dual regulation highlights ZNF281 as a potential driver of EMT and tumor progression in prostate cancer (**Fig. 4I**).

### Generation of the orally bioavailable ZNF281 Interfering Molecule (Oral ZIM)

In our previous study(13), the early version of ZIM was shown to directly disrupt ZNF281-DNA binding, and this effect was abolished by mutations within the predicted binding interface (F326A/I327A), supporting its on-target engagement. Building on this validated framework, we developed a structurally optimized ZIM containing fewer hydrophobic fluorinated groups. This optimized ZIM supported oral dosing in preclinical *in vivo* studies while retaining its biological activity against ZNF281 (**Fig. 5A, B**, **Supplementary Fig. S5A**). Electrophoretic mobility shift assay confirmed that oral ZIM reduced ZNF281-DNA interaction (**Fig. 5C**). The oral version ZIM was used throughout this study to evaluate its translational potential. Notably, ChIP-seq metagene and heatmap analyses revealed strong enrichment of ZNF281 at promoter regions (± 2 kb from transcription start site, TSS) in LNCaP cells under vehicle conditions. Upon oral ZIM treatment, ZNF281 occupancy at promoter-proximal regions was markedly reduced, indicating that ZIM globally impairs ZNF281 binding at gene promoters (**Fig. 5D**). A similar reduction of ZNF281 occupancy at promoter-proximal regions was also observed in ZIM-treated DU145 cells (**Supplementary Fig. S5B**). Together, these data demonstrate that oral ZIM could disrupt ZNF281 binding at gene promoters, leading to transcriptional changes of its direct target genes.

**Figure 5.**
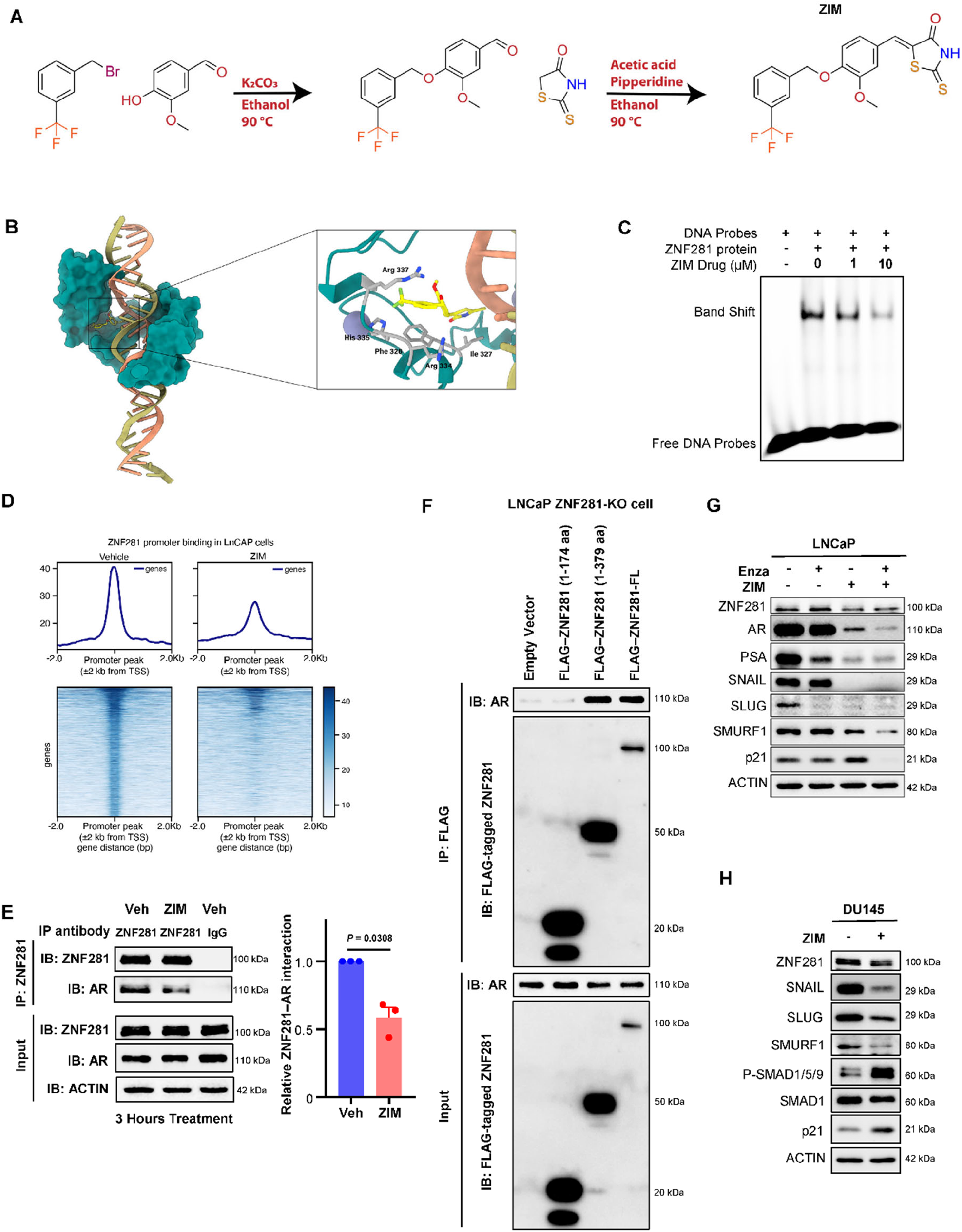
Generation of the orally bioavailable ZNF281 Interfering Molecule (Oral ZIM). **A,** Reaction scheme illustrating the full synthetic sequence of ZIM. Vanillin was O-alkylated with 3- (trifluoromethyl)benzyl bromide under basic conditions (K2CO3, EtOH, heat) to afford int-ZIM (4-((3-(trifluoromethyl)benzyl)oxy)-3-methoxybenzaldehyde) as the key benzylated aldehyde intermediate. Int-ZIM was subsequently converted to ZIM ((5Z)-5-(4-((3- (trifluoromethyl)benzyl)oxy)-3-methoxybenzylidene)-2-thioxo-1,3-thiazolidin-4-one) via a rhodanine-based Knoeven.agel-type condensation (EtOH, catalytic piperidine, AcOH, heat), providing the corresponding Z isomer. **B,** Predicted binding mode of ZIM at the ZNF281 DNA binding interface. The left view shows the overall ZNF281–DNA assembly showing the zinc-finger region wrapped around the DNA. The right view is a zoomed-in representation of the ZIM docked into ZNF281 DBD binding pocket. In this pose, ZIM is predicted to engage several residues (326, 327, 334, 335 and 337) on the ZNF281/DNA interface and to sterically occlude, and/or disrupt, key protein–DNA contacts, thereby interfering with ZNF281 association with DNA. **C,** Electrophoretic mobility shift assay (EMSA) showing that ZIM disrupts ZNF281–DNA probe binding, as indicated by decreased band-shift signals. **D,** Metaplot (top) and heatmap (bottom) showing ZNF281 ChIP-seq signal centered on promoter regions (±2 kb from transcription start sites, TSS) in LNCaP cells under vehicle or ZIM treatment. Each row represents a gene promoter, ranked by ZNF281 signal intensity. **E,** Representative western blot and quantification of co-immunoprecipitation analysis in LNCaP cells treated with ZIM (10 μM) for 3 hours. Co-immunoprecipitated AR was normalized to immunoprecipitated ZNF281, showing that ZIM reduced the ZNF281–AR interaction before detectable changes in the protein abundance of either ZNF281 or AR. **F,** Co-immunoprecipitation analysis of FLAG-tagged full-length (FL) and truncated ZNF281 constructs in ZNF281-knockout LNCaP cells. Immunoprecipitation with anti-FLAG followed by immunoblotting for AR showed that the ZNF281 region spanning amino acids 175–379 was required for its association with AR. Expression and immunoprecipitation of the FLAG-tagged constructs were confirmed by anti-FLAG immunoblotting. **G,** Western blot analysis of LNCaP treated with ZIM (10 µM) and enzalutamide (10 µM) for 48 hours. **H,** Western blot analysis of DU145 treated with ZIM (10 µM) for 48 hours. Graphs show the mean ± SEM from three independent biological replicates. Statistical significance was determined using t test with Welch’s correction.

Co-immunoprecipitation assays further demonstrated that ZIM reduced the association between ZNF281 and AR before detectable changes in the abundance of either protein **(Fig. 5E)**. Notably, Nicolai et al. identified the zinc-finger domain within the amino acid 175–379 region as being required for both ZNF281 recruitment to damaged DNA and its interactions with the non-homologous end-joining proteins. Disruption of this region impaired both the DNA-binding and protein-interaction activities of ZNF281(23). Consistent with these findings, co-immunoprecipitation assays using truncated ZNF281 constructs showed that the amino acid 175– 379 region was also required for the association between ZNF281 and AR (**Fig. 5F**). Together, these findings suggest that this region may constitute a multifunctional interface involved in both DNA binding and protein-complex formation. ZIM may therefore inhibit both activities by perturbing this interface.

Western blot analysis revealed that ZIM downregulated AR and its downstream target (PSA) in LNCaP cells (**Fig. 5G, Supplementary Fig. S6A**). The AR-suppressive effect of ZIM-mediated ZNF281 inhibition exceeded that of enzalutamide (a first-line treatment for metastatic castration-sensitive and castration-resistant prostate cancers). In both LNCaP and DU145 cell lines, ZIM lowered EMT-associated markers SNAIL and SLUG (**Fig. 5G, H, Supplementary Fig. S6A, B**). Moreover, ZIM downregulated SMURF1, and increased p-SMAD1/5/9 and p21 expression, indicating its potential antiproliferation effect on prostate cancer (**Fig. 5G, H, Supplementary Fig. S6A, B**). Additionally, in PC3 cells, ZIM treatment reduced SMURF1 and SNAIL expression while increasing the p-SMAD1/5/9-to-total SMAD1 ratio and p21 levels (**Supplementary Fig. S6C**). These findings recapitulated the effects of genetic ZNF281 suppression on the SMURF1-associated proliferative and SNAIL-associated EMT pathways.

ZIM was sufficient to inhibit ZNF281-DNA binding in EMSA, reduce the abundance of ZNF281-regulated proteins, and inhibit the proliferation of parental LNCaP cells, whereas its antiproliferative effect was attenuated in ZNF281-KO cells (**Fig. 5C and Supplementary Fig. S6D, E**). To assess its potential off-target activity, ZIM was evaluated across a panel of 44 pharmacologically relevant targets, including GPCRs, ion channels, transporters, kinases, and nuclear receptors (InVEST44 Report, **Supplementary Table S2**). ZIM exhibited minimal off-target interactions, with most targets showing activity values within 75-125% or around 0% (agonist model) of baseline, indicating negligible modulation. Importantly, no significant inhibition was observed for key safety-related targets, including the hERG potassium channel, major neurotransmitter receptors, and transporters.

Consistent with its proposed mechanism, ZIM showed only weak activity toward AR, suggesting that its modulation of AR signaling is unlikely to arise from direct receptor binding. Together, these results demonstrate that ZIM possesses a favorable selectivity profile with minimal off-target activity, supporting its function as a ZNF281-interfering molecule.

### ZIM inhibited cancer growth and metastasis *in vitro* and in prostate cancer preclinical models

The antitumor efficacy of ZIM was first evaluated *in vitro* using cell lines. CCK-8 assays showed that ZIM significantly decreased cell viability, with greater inhibitory effects than enzalutamide (**Fig. 6A**). No significant synergistic effect was observed for ZIM combined with enzalutamide in either cell line. ZIM also markedly reduced transwell migration (**Fig. 6B**) and invasion (**Fig. 6C**), again outperforming enzalutamide. Consistent inhibitory effects on proliferation, migration, and invasion were also observed in PC3 cells, extending the antitumor activity of ZIM to an additional AR-negative model (**Supplementary Fig. S7A-C**).

**Figure 6.**
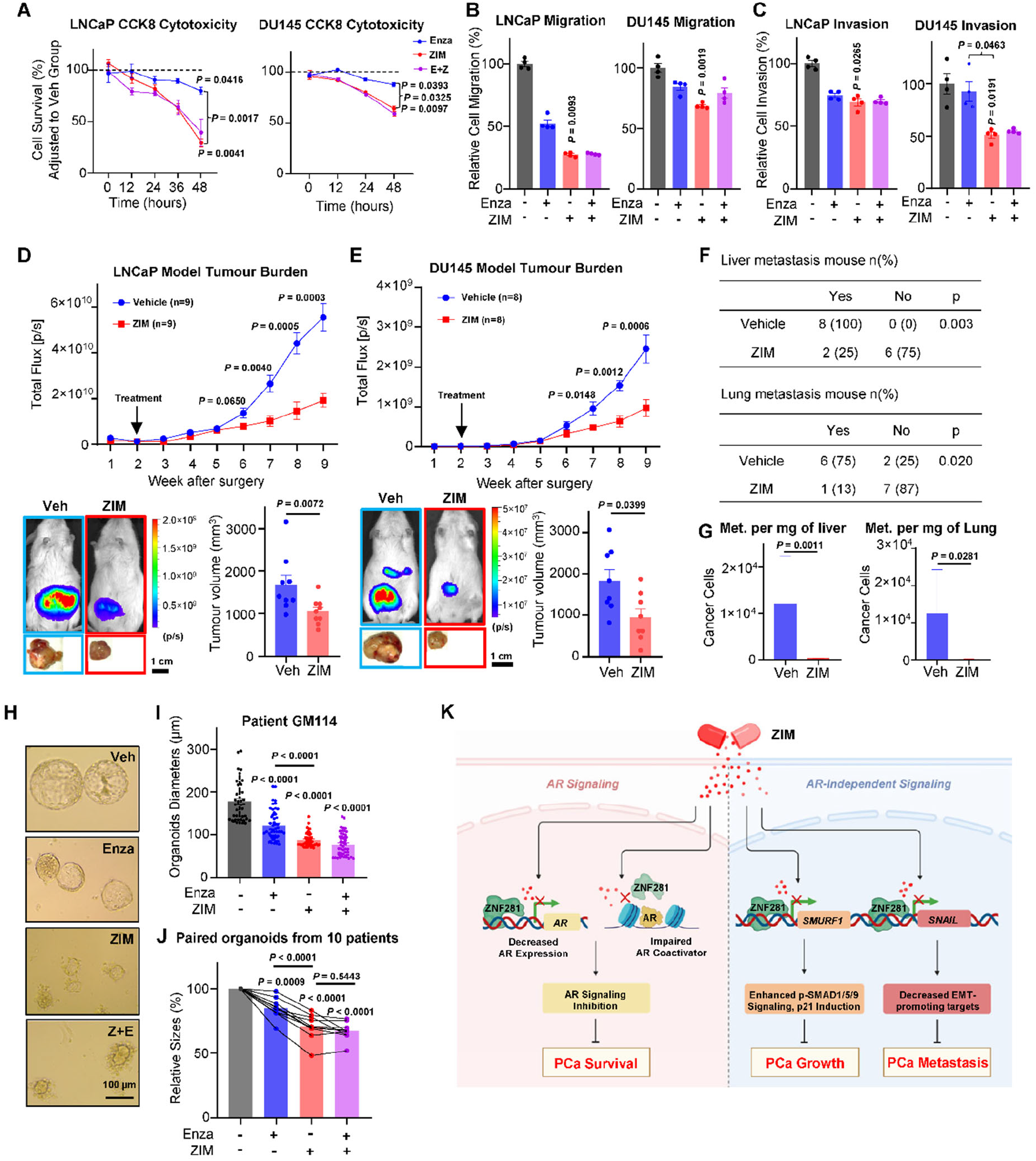
ZIM inhibited cancer growth and metastasis *in vitro* and in prostate cancer preclinical models. **A,** *In vitro* CCK-8 cell proliferation assay showing the effects of ZIM (10 µM), enzalutamide (10 µM) and combination treatment on cell proliferation in LNCaP and DU145 cells. **B,** Transwell migration assay in LNCaP and DU145 showing that ZIM (10 µM) treatment inhibited cancer cell migration. **C,** Transwell invasion assay showing that ZIM (10 µM) treated LNCaP and DU145 cells exhibited reduced invasive capacity. **D,** Statistical data and representative bioluminescence images of ZIM treatment in an orthotopic LNCaP xenograft mouse model. Tumor growth was monitored weekly by *in vivo* bioluminescence imaging. LNCaP tumors treated with ZIM (250 mg/kg, oral gavage, daily for 7 weeks) grew significantly more slowly and exhibited smaller endpoint tumor volumes. **E,** Statistical data and representative bioluminescence images of ZIM treatment on the orthotopic DU145 xenograft mouse model. DU145 tumors treated with ZIM (250 mg/kg, oral gavage, daily for 7 weeks) grew significantly more slowly, exhibited smaller endpoint tumor volumes, and showed fewer metastatic lesions. **F,** Statistical analysis of mice with liver or lung metastasis. **G,** Quantification of tumor burden in liver or lung metastases. **H,** Representative images and statistical data of patient-derived organoids GM114 treated with ZIM (20 µM), enzalutamide (20 µM), or combination treatment for 7 days. **I,** Statistical data of patient-derived organoids from patient GM114 treated with ZIM (20 µM), enzalutamide (20 µM), or combination treatment for 7 days. **J,** Statistical data of patient-derived organoids from ten patients treated with ZIM (20 µM), enzalutamide (20 µM), or combination treatment for 7 days. **K,** Schematic illustration of the mechanisms of action of ZIM in prostate cancer. Graphs show the mean ± SEM from independent biological replicates. Statistical significance was determined using RM one-way ANOVA with Bonferroni post hoc multiple-comparison test (**A, J**), Kruskal-Wallis test with Dunn’s post hoc multiple comparisons test (**B, C**), Mann–Whitney test (**D, E, G**), one-sided Fisher’s exact test (**F**), one-way ANOVA with Bonferroni post hoc multiple-comparison test (**I**).

The therapeutic potential of oral ZIM was evaluated *in vivo* using orthotopic xenograft models. The *in vivo* dose-selection strategy was informed by the biologically active concentration identified *in vitro*. Pharmacokinetic analysis showed that a single oral dose of ZIM at 250 mg/kg resulted in measurable systemic exposure, with an estimated oral bioavailability (*F*) of 10.6% (**Supplementary Fig. S8 and Supplementary Table S3**). In the ZIM safety study, healthy, non-tumor-bearing mice, ZIM was well tolerated, with no notable toxicity observed. Liver and kidney function remained within normal ranges, blood cell counts and body weight were maintained after seven weeks of daily treatment (**Supplementary Table S4, Supplementary Fig. S9**). Treatment of tumor mice began two weeks post-tumor implantation with the same regimen. HPLC-MS/MS analysis showed gastrointestinal absorption and systemic distribution of oral ZIM, including its accumulation within prostate tumors (**Supplementary Table S5**). After seven weeks of treatment, ZIM markedly reduced tumor luminescence intensity and endpoint tumor volume in both LNCaP and DU145 models (**Fig. 6D, E**). In DU145-bearing mice, ZIM significantly decreased the ratio of mice with liver metastasis (100% to 25%) and lung metastasis (75% to 13%, **Fig. 6F**). It also reduced the tumor burden at the metastatic sites (**Fig. 6G**).

To examine the effects of ZIM on ZNF281-associated signaling *in vivo*, tumor lysates were analyzed by western blot. ZIM-treated LNCaP tumors showed variable reductions in ZNF281 and its downstream proteins, including AR, PSA, SNAIL and SMURF1, whereas reductions in SNAIL and SMURF1 were observed in ZIM-treated DU145 tumors, compared to their vehicle groups (**Supplementary Fig. S10 A, B)**. These findings suggest that ZIM can suppress both AR-dependent and AR-independent ZNF281-regulated pathways *in vivo*.

To assess its clinical relevance, ZIM was tested in patient-derived organoid (PDO) models established from treatment-naïve primary prostate tumor specimens obtained from ten different patients (**Supplementary Fig. S11, Supplementary Table S6**). ZIM inhibited organoid growth in a dose-dependent manner **(Supplementary Fig. S12)**. Organoids were further treated with vehicle, enzalutamide, ZIM, or the combination of ZIM and enzalutamide (**Fig. 6H**). ZIM significantly inhibited organoid growth in all ten cases, and showed superior efficacy compared to enzalutamide (**Fig. 6I, J**). Likely due to effective suppression of AR signaling by ZIM alone, combination treatment did not yield a significant synergistic benefit.

Collectively, the mechanisms of action of ZIM on metastatic prostate cancer (**Fig. 6K**) include, <u>1.</u> Inhibition of AR signaling by repressing AR expression and AR transcriptional activity; <u>2.</u> The AR-independent mechanisms involving inhibition of SMURF1-mediated tumor growth and SNAIL-mediated metastasis.

## DISCUSSION

While AR signaling remains a central oncogenic axis in prostate cancer, accumulating evidence indicates that AR pathway inhibition alone is insufficient to control late-stage disease, particularly in the context of AR-independent growth and metastasis(24–26). In this study, we have found that ZNF281 could be a novel therapeutic target for metastatic prostate cancer. It regulated AR signals by promoting AR transcription and acting as a coactivator to facilitate AR transcriptional activity. Furthermore, it also promoted tumor growth, progression, and metastasis through distinct AR-independent tumor-promoting pathways. As the first-in-class inhibitor of ZNF281, the oral ZIM effectively inhibited primary tumor growth and metastasis of prostate cancer *in vitro*, *in vivo*, and in patient-derived organoid models. It simultaneously targeted metastatic castration-sensitive and castration-resistant prostate cancer and may therefore represent an effective approach to limit disease progression and cancer-specific mortality.

Unlike classical therapy targeting the AR itself, ZNF281 is an upstream regulator that directly promotes AR expression. Importantly, ZNF281 depletion impaired AR signaling even under androgen-depleted conditions, suggesting that ZNF281 contributed to ligand-independent AR activation, which is a hallmark of CRPC.

Moreover, targeting AR coactivator networks has been acknowledged as a strategy to overcome ADT resistance, including AR-ligand independence(27). ZNF281 was previously identified as an AR-interacted protein but lacked functional characterization in prostate cancer(14). Our study revealed that ZNF281 may function as a facilitator that promotes AR transcriptional activity. This mechanistic role may explain how its inhibition led to a more profound suppression of AR signaling than enzalutamide in our models.

A major limitation of AR-targeted therapies is their ineffectiveness against AR-low or AR-negative prostate cancer that frequently emerge under prolonged AR pathway suppression(28–30). Notably, ZNF281 promotes tumor growth, invasion, and metastasis in AR-negative DU145 and PC3 cells, demonstrating that its oncogenic role extends beyond AR signaling.

Our study identified SMURF1 as a direct downstream target of ZNF281 in both AR-positive and AR-negative prostate cancer cells. Although SMURF1 has been reported as an AR-regulated gene that enhances prostate cancer migration and invasion(31), our findings suggested that, within the ZNF281-regulated network, SMURF1 primarily mediated tumor cell proliferation and was not a major contributor to migration or invasion in LNCaP and DU145 cells. Instead, these invasive phenotypes appear to be mediated predominantly through the SNAIL-associated EMT program, consistent with the established role of ZNF281 in activating EMT(21,22,32). Moreover, ZNF281 sustains SMURF1 expression in the absence of AR, providing a mechanism for maintaining proliferative signaling in AR-negative prostate cancer. Thus, by regulating SMURF1-associated proliferative axis and the SNAIL-associated EMT program in parallel, ZNF281 may coordinate tumor growth and metastasis progression across different AR contexts of prostate cancer.

Although a recent study identified ZNF281 as a transcriptional repressor of GPX4 and SLC3A2 that promotes ferroptosis in prostate cancer(33), our findings reveal a distinct role for ZNF281 as a regulator of AR signaling and metastatic progression. These findings are not necessarily mutually exclusive, as the two studies address different biological processes of prostate cancer with distinct downstream pathways regulated by ZNF281.

These mechanisms are consistent with the therapeutic effects of ZIM, which exhibited a robust anti-AR effect and retained therapeutic efficacy in the AR-negative PCa models. The absence of synergistic effects between ZIM and enzalutamide suggests that ZIM alone may already maximally suppressed AR transcriptional output in LNCaP cells. Importantly, ZIM demonstrated robust efficacy in patient-derived organoid models, which preserve interpatient heterogeneity and clinically relevant features of prostate cancer(34,35). Moreover, the conserved molecular and functional effects observed in LNCaP, DU145 and PC3 cells support the activity of ZNF281 inhibition across distinct metastatic prostate cancer contexts.

Nevertheless, these models do not fully capture the heterogeneity of metastatic CRPC, particularly treatment-emergent, lineage-plastic and neuroendocrine-like states(28,36). Defining the role of ZNF281 in lineage plasticity, neuroendocrine differentiation and therapeutic resistance using advanced model systems, including larger, molecularly annotated PDO cohorts, represents an important direction for future investigation. These studies may also help identify ZNF281-based molecular features associated with ZIM sensitivity and inform future patient selection strategies. Furthermore, the AR-independent mechanism of ZNF281 suggests that ZIM may have therapeutic potential across a broad spectrum of cancers, warranting further investigation.

## METHODS

### Ethics Statement

All animal studies were approved by the University of Alberta Animal Care and Use Committee (ACUC; AUP00004359) and conducted in accordance with institutional and national guidelines. Animals were monitored daily by animal facility staff as part of routine husbandry and three times per week by study investigators. Animals displaying signs of discomfort or pain, including weight loss, decreased respiration, or impaired mobility as defined by humane endpoints, were euthanized. All data were included in the analysis except when morbidity or mortality was attributable to surgical procedures or oral gavage. Investigators involved in data collection and analysis were blinded to group allocation.

### Orthotopic Xenograft Model of Human Prostate Cancer

The orthotopic xenograft of prostate cancer model was performed as previously described(37). Specifically, 1 × 10^6^ of luciferase-stably expressed DU145 or LNCaP cells were suspended in 20 μL of injection solution consisting of 50% growth medium and 50% Matrigel (Corning Matrigel, CLS354234). Cell suspension was placed on ice and injected within 30 mins to ensure viability. The NSG male mice (NOD.Cg-Prkdc^scid^ Il2rg^tm1Wjl^/SzJ, Strain No. 005557, The Jackson Laboratory) were provided by Dr. Lynne Postovit’s lab. All 6-to 10-week-old SCID mice were randomly separated into each experimental group without prior designation. General anesthesia was followed by an open abdominal surgery to locate the mouse prostate glands. The human prostate cancer cells suspended solution was injected into the left anterior lobe of the prostate gland. It generally takes two weeks for the tumor to be stably established in the mouse anterior prostate. The whole study lasted for 9 weeks after the cancer cell injection. In the drug-treated experiments, ZIM (250 mg/kg) or vehicle (10% DMSO in corn oil) was given daily by gavage feeding two weeks after the surgery and lasted for seven weeks.

### Live Animal Imaging

Human prostate cancer cell lines stably expressing luciferase were used. Tumor growth and metastatic dissemination were monitored weekly by *in vivo* bioluminescence imaging (IVIS Spectrum) following intraperitoneal injection of D-luciferin (GoldBio, 115144-35-9), according to the manufacturer’s instructions, until study endpoints were reached.

### Quantification of Mice Metastasis

Distal organs, including liver and lung of the tumor-bearing sacrificed mice, were harvested, washed with PBS, and fixed with enough RNA*later*™ Stabilization Solution (Thermo Fisher Scientific, AM7020). Tissues were pulverized and weighed 30 mg for further RNA isolation and RT-qPCR experiments.

Cancer cell-specific primers human B2M (Thermo Fisher Scientific, Hs00187842_m1) and Luciferase (Thermo Fisher Scientific, Mr03987587_mr) were used to quantify the tumor cell burdens in tissue, while the mouse B2M (Thermo Fisher Scientific, Mm00437762_m1) was used as the internal control. Only tissue samples with both positive human B2M and luciferase expression (compared to tissues from healthy mice) were considered positive for organ metastasis.

To quantify tumor burden within metastatic organs, standard curves were generated using defined numbers of luciferase-labeled DU145 cells mixed with 30 mg of normal lung or liver tissue. RNA was isolated, and RT–qPCR was performed using the same protocol as for tumor-bearing mouse organs. Relative expression levels of human B2M were calculated using the 2^-ΔCT^ method and used to estimate tumor burden (cancer cells number per mg of tissue).

### IF, H&E Staining

Paired tissue sections from 15 patients who underwent radical prostatectomy with bilateral lymph node dissection were obtained from the APCaRI (Alberta Prostate Cancer Research Initiative) biobank. Regions of interest (primary tumor lesions and metastatic lesions) were identified and circled by a pathologist based on pathological evaluation. Immunofluorescence staining was subsequently performed, followed by imaging using a Zeiss LSM 900 confocal laser-scanning microscope. The following primary antibodies were used for IF staining at a dilution of 1:100: ZNF281 (Santa Cruz Biotechnology, sc-166933), ZNF281 (Sigma Aldrich, HPA051228), AR (Cell Signaling Technology, 5153S), PSA (Cell Signaling Technology, 5365), AMACR (Cell Signaling Technology, 29256), CD44 (Cell Signaling Technology, 5640), and Keratin8/18 (Cell Signaling Technology, 51274). The following secondary antibodies were used for IF staining at a dilution of 1:1000: donkey anti-mouse IgG (H+L) Alexa Fluor 647 secondary antibody (Thermo Fisher Scientific, A-31571), donkey anti-mouse IgG (H+L) Alexa Fluor 546 secondary antibody (Thermo Fisher Scientific, A10036), and donkey anti-rabbit IgG (H+L) Alexa Fluor 647 secondary antibody (Thermo Fisher Scientific, A-31573). Zeiss ZEN Blue software was used for quantification of image signal intensity.

Liver and lung tissue of the xenografted mice were harvested, washed with cold PBS, and immediately fixed in 10% neutral-buffered formalin (Sigma-Aldrich, HT501320) for 24 hours. Fixed tissues were then processed and subjected to hematoxylin and eosin (H&E) staining by the University of Alberta LMP Pathology Core.

### Cell Lines and Cell Culture

Human Prostate cell lines LNCaP clone FGC (ATCC, CRL-1740), DU145 (ATCC, HTB-81), PC3 (ATCC, CRL-1435), HPrEC (ATCC, PCS-440-010) and RWPE1 (ATCC, CRL-3607) were obtained from the American Type Culture Collection (ATCC, USA) and cultured according to the manufacturer’s instructions. The luciferase/GFP dual-labeled DU145 (GeneCopoeia, SLC023) and luciferase/GFP dual-labeled LNCaP (FenicsBio, CL-1538) cell lines were used in the *in vivo* experiments. LNCaP and DU145 ZNF281-KO cell lines were generated from GenScript. All cancer cell lines were cultured in medium containing 10% FBS, 1% Penicillin-Streptomycin-Amphotericin B (Gibco, 15240062), Plasmocin Prophylactic 5ug/ml (InvivoGen, ant-mpp) with 5% CO_2_ conditions. All cells were routinely tested for mycoplasma using the LookOut Mycoplasma PCR Detection Kit (Sigma-Aldrich, MP0035) every month.

### *In Vitro* Cell Proliferation, Migration, and Invasion Assays

Cancer cell proliferation and drug-induced cytotoxicity were assessed using the Cell Counting Kit-8 (CCK-8; Sigma-Aldrich, 96992). Cells were synchronized by overnight incubation in serum-free medium prior to experimentation. A total of 5,000 cells were seeded in 96-well plates in 100 μL of complete medium. Subsequently, 10 μL of CCK-8 solution was added to each well, and plates were incubated for 1.5 hours at 37°C in a humidified incubator with 5% CO₂ before absorbance was measured using a microplate reader.

Transwell migration and invasion assays were performed using 96-well plates with Boyden chambers (migration: 5.0 μm pore size; Abcam, ab235693; invasion: 8.0 μm pore size chambers coated with basement membrane extract; Abcam, ab235697) according to the manufacturer’s instructions. Cells were synchronized overnight in serum-free medium before seeding. A total of 20,000 cells suspended in 100 μL of serum-free medium were added to the upper chamber, while 200 μL of medium containing 10% FBS was added to the lower chamber as a chemoattractant. Plates were incubated at 37°C for 48 hours.

### *In Vitro* Cell Transfection

Small interfering RNA (siRNA): Cells were grown in 6-well plate to 50-60 % confluence and then transfected with 30 pmol siRNA diluted in Opti-MEM medium and mixed with Lipofectamine RNAiMAX reagent (Invitrogen, 13778075). The mixture was incubated for 10 minutes at room temperature and then added to the cells for 72 hours. siRNA used in the study: siZNF281 (Santa Cruz Biotechnology, sc-88283), siAR (Thermo Fisher Scientific, s1538). Adenoviral infection: Adenoviruses were used at a final concentration of 500 multiplicity of infection overnight in antibiotic-free media, and then changed to fresh complete media for 48 to 72 hours.

### Protein Extraction and Western Blot

Cell pellets were lysed in RIPA buffer (Thermo Fisher Scientific, 89900) supplemented with 1 mmol/L PMSF (Sigma-Aldrich, 93482) and 1× protease and phosphatase inhibitor cocktails (Sigma-Aldrich, P2714). Lysates were incubated on ice for 25 minutes and clarified by centrifugation at 12,000 × g for 15 minutes at 4°C. Protein concentrations were determined using a BCA protein assay kit (Thermo Fisher Scientific, 23209). The protein lysates were eluted with an equal amount of 2× Laemmli Sample buffer (Sigma Aldrich S3410) and boiled for 5 minutes at 100°C. Equal amounts of protein were resolved by SDS-PAGE and transferred onto 0.45μm nitrocellulose membranes. Membranes were blocked with 5% silk milk for 1 hour at room temperature and then incubated overnight at 4°C with diluted primary antibodies (1:1000). After washing with TBST (Tris-buffered saline containing 0.1% Tween 20), membranes were incubated with horseradish peroxidase-coupled secondary antibodies (1:3000) in TBST at room temperature for 1 hour. Protein signals were detected using enhanced chemiluminescence (ECL) reagents (Bio-Rad, 1705062) and visualized with the ChemiDoc™ MP Imaging System (Bio-Rad, 12003154). Band intensities were quantified using ImageJ software and normalized to loading controls. The following primary antibodies were used in western blot: ZNF281 (Santa Cruz Biotechnology, sc-166933), ZNF281 (Sigma Aldrich, HPA051228), AR (Abcam, ab52615), PSA (Proteintech, 10679-1-AP), TMPRSS2 (Santa Cruz Biotechnology, sc-515727), p300 (Cell Signaling Technology, 86377), Acetyl-H3K27 (Cell Signaling Technology, 8173), H3 (Cell Signaling Technology, 14269), SNAIL (Cell Signaling Technology, 3879), SLUG (Cell Signaling Technology, 9585), E-Cadherin (Cell Signaling Technology, 3195), SMURF1 (Cell Signaling Technology, 2174), p-SMAD1/5/8 (Cell Signaling Technology, 13820), SMAD1 (Cell Signaling Technology, 6944), p21 (Cell Signaling Technology, 2947), ZNF217 (Cell Signaling Technology, 82306), ACTIN (Abcam, ab179467).

### Coimmunoprecipitation

Target cells were lysed using Pierce™ IP Lysis Buffer (Thermo Fisher Scientific, 87788) supplemented with 1 mmol/L PMSF and 1× protease and phosphatase inhibitor cocktails. Total protein concentration was determined using a BCA assay. While 50 μg of protein lysate was reserved as the input control, 400 μg of protein was incubated with 4 μg of ZNF281 antibody, and an equal amount of protein lysate was incubated with 4 μg of IgG isotype control antibody (Cell Signaling Technology, 5415). The antibody-protein complexes were incubated with rotation overnight at 4 °C. Subsequently, Dynabeads™ Protein A (Invitrogen, 10008D) were added and incubated at 4 °C for 2 hours. The beads were then separated, washed, and eluted with an equal volume of IP lysis buffer and 2× Laemmli sample buffer, followed by boiling at 100 °C for 5 minutes.

### RNA Extraction and RT-qPCR

Total RNA was isolated using QIAzol Lysis Reagent (Qiagen, 79306) according to the manufacturer’s instructions. Quantitative reverse transcription PCR (qRT-PCR) was performed using the TaqMan™ RNA-to-CT™ 1-Step Kit (Applied Biosystems™, 4392938) on a QuantStudio™ 7 Flex Real-Time PCR System (Thermo Fisher Scientific, 4485701), following the manufacturer’s protocols. The following TaqMan® Gene Expression Assays were purchased from Thermo Fisher Scientific and used in this study: ZNF281 (Hs00273550_s1), AR (Hs00171172_m1), PSA (Hs03063374_m1), KLK2 (Hs00428383_m1), TMPRSS2 (Hs05024838_m1), EP300 (Hs00914223_m1), B2M (Hs00187842_m1), Mouse B2M (Mm00437762_m1), Luciferase (Mr03987587_mr).

### RNA Sequencing and Analysis

Raw RNA-sequencing reads were quality assessed using FastQC (v0.12.1) and preprocessed with Fastp (v1.0.0)(38) for adapter removal and quality filtering (Q < 20). Reads were aligned to the human reference genome (GRCh38.p14) using STAR (v2.7.11b)(39) in two-pass mode with GENCODE v46 annotation. Gene-level counts were generated using HTSeq-count (v2.0.5)(40) with stranded settings. Differential expression analysis was performed in R (v4.4.2)(41) using the limma-voom framework(42–44) with TMM normalization and empirical Bayes moderation (robust = TRUE). Genes with FDR ≤ 0.05 and |log₂ fold change| ≥ 0.6 were considered significant. Functional enrichment analyses were conducted using clusterProfiler for Gene Ontology and KEGG pathways, ReactomePA(45) for Reactome pathways, and GSEA(46) using pre-ranked log₂ fold-change values. Signaling pathway activity was inferred using PROGENy(47) implemented via decoupleR(48). Analyses were performed independently for LNCaP and DU145 cells. Data visualization was performed using ggplot2, pheatmap, and patchwork. Read preprocessing and alignment were conducted using Galaxy (v25.0)(49), with downstream analyses performed in R.

### ChIP and ChIP-qPCR

ChIP was performed using the SimpleChIP® Plus Enzymatic Chromatin IP Kit (Cell Signaling Technology, 9005) with ChIP-grade antibodies against ZNF281 (Santa Cruz Biotechnology, sc-166933 X) and AR (Cell Signaling Technology, 5153S), according to the manufacturer’s instructions. The immunoprecipitated DNA was amplified using Fast SYBR™ Green Master Mix (Applied Biosystems™, 4385612) on a QuantStudio™ 7 Flex Real-Time PCR System. The DNA primers used in this study include:

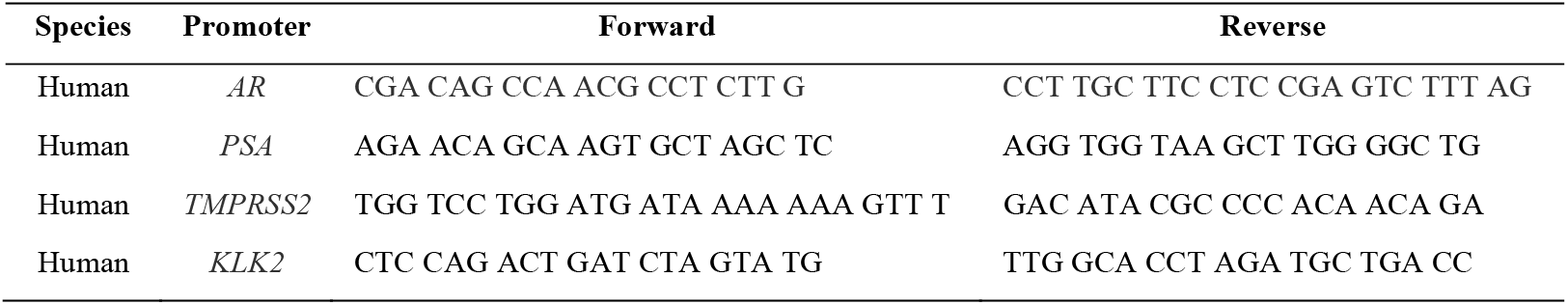

### ChIP Sequencing and Analysis

ChIP-seq libraries were sequenced using paired-end 2 × 101 bp reads. Raw reads were processed using fastp (v0.23.2) for adapter trimming and quality filtering, and aligned to the human reference genome (hg38) using Bowtie2(50) with default parameters. Aligned reads were sorted and indexed using SAMtools (v1.18). Before merging, concordance among biological replicates was assessed by calculating pairwise Pearson correlation coefficients based on genome-wide ChIP-seq read coverage using deepTools. Unsupervised hierarchical clustering was performed based on the resulting correlation matrices. Biological replicates were merged only after their reproducibility had been confirmed. Fragment length distributions were assessed using deepTools to confirm library quality and enrichment. Peak calling was performed using MACS2 (v2.1.2)(51) with paired-end support and matched input controls using the human genome size parameter. Peaks were annotated using GENCODE v46 gene annotations, with promoter regions defined as ±2 kb relative to TSS. Signal enrichment across promoter regions was analyzed using deepTools to generate normalized coverage matrices, metaplots, and heatmaps. For locus-specific visualization, normalized ChIP-seq signal tracks were visualized using SparK to generate IGV-style coverage plots(52).

### Drug Design and Discovery

ZNF281 was modeled by homology modeling using SWISS-MODEL, with the mouse ZFP568-ZnF1-11-DNA complex as the structural template (sequence similarity 34.51%)(53). The resulting model was prepared and optimized in the Maestro Schrödinger suite (Schrödinger Release 2024-1: Maestro, Schrödinger, LLC, New York, NY, 2024) using the Protein Preparation Wizard, including completion of missing side chains and assignment of protonation states with Epik(54).

To identify a starting chemical scaffold, an in-house compound collection was subjected to high-throughput screening, which led to selection of the ZIM backbone for subsequent structure-based optimization. For pose generation, blind docking of ZIM into the putative ZNF281 pocket was performed with VinaMPI using a 30 × 30 × 30 Å³ grid and 0.375 Å spacing(55). Docking hits were subsequently refined using Schrödinger Ligand Designer and predicted physicochemical and pharmacokinetic properties were assessed using QikProp and SwissADME(56). Graphical representations were generated with ChimeraX(57).

### Synthesis of (5Z)-5-(4-((3-(trifluoromethyl)benzyl)oxy)-3-methoxybenzylidene)-2-thioxo-1,3-thiazolidin-4-one (ZIM)

4-((3-trifluoromethyl)benzyl)oxy)-3-methoxybenzaldehyde (Int-ZIM): Vanillin (1.0 equiv) 3- (trifluoromethyl)benzyl bromide (1.0 equiv) and potassium carbonate (1.0 equiv) were combined in ethanol (50 mL) in a round-bottom flask, and the mixture was heated at 90 °C for 12 hours with magnetic stirring. Reaction progress was followed by TLC. After completion, the product precipitated as a white crystalline solid. The solid was collected by filtration and washed sequentially with water (10 mL) and ice-cold ethanol (10 mL) to afford Int-ZIM. ^1^H NMR (600 MHz, CDCl3) δ 9.88 (s, 1H), 7.74 (s, 1H), 7.66 (d, J = 7.6 Hz, 1H), 7.62 (d, J = 7.8 Hz, 1H), 7.54 (t, J = 7.7 Hz, 1H), 7.48 (m, 1H), 7.41 (m, 1H), 7.01 (d, J = 8.2 Hz, 1H), 5.29 (s, 2H), 3.97 (s, 3H).

Int-ZIM (1.0 equiv) and rhodanine (1.2 equiv) were placed in a round-bottom flask, followed by addition of a catalytic amount of piperidine and acetic acid. Ethanol (50 mL) was then added, and the reaction mixture was heated at 90 °C for 12 hours with magnetic stirring. After cooling to room temperature, a bright yellow solid formed and was isolated by filtration. The crude product was washed with cold ethanol (30 mL) and water (30 mL) to give ZIM. The product was assigned as the Z isomer based on the reported methine proton chemical shift(58). ^1^H NMR (600 MHz, DMSO – d6) δ 13.78 (s, 1H), 7.84 (s, 1H), 7.78 (d, J = 7.80 Hz, 1H), 7.74 (d, J = 7.77 Hz, 1H), 7.67 (t, J = 7.7 Hz, 1H), 7.62 (s, 1H), 7.27 – 7.22 (m, 2H), 7.20 (dd, J = 8.5, 2.1 Hz, 1H), 5.30 (s, 2H), 3.86 (s, 3H) (**Supplementary Fig. S13A**). ^13^C NMR (600 MHz, DMSO – d6) δ 195.93, 169.84, 150.34, 149.79, 138.50, 132.56, 132.33, 130.10, 129.79, 127.34, 125.53, 125.24, 124.79, 124.26, 114.26, 114.22, 69.58, 56.13. (**Supplementary Fig. S13B**). High-resolution full scan mass spectrometry in electrospray negative mode was performed, giving the ion at m/z 424.0298 corresponding to the chemical formula C19H14F3NO3S2 (theoretical negative m/z 424.0294) on Q Exactive Orbitrap Elite Mass spectrometer (Thermo Scientific, San Jose, CA) (**Supplementary Fig. S13C**).

### Pharmacokinetic Study

The pharmacokinetic properties of ZIM were evaluated by Ichor Life Sciences in male C57BL/6 mice aged 6–8 weeks. Mice received a single oral dose of ZIM at 250 mg/ kg or a single intravenous dose at 2.5 mg/ kg (15 mice per administration route). For oral administration, ZIM was formulated in 10% DMSO and 90% corn oil (v/v) and administered by oral gavage. For intravenous administration, ZIM was formulated at 1.25 mg/ ml in 10% N-methyl-2-pyrrolidone, 5% Cremophor EL and 85% sterile water (v/v/v) and administered by tail-vein injection.

Blood samples were collected at 15, 30, 60, 120, 240, 360, 480 and 1,440 min after oral administration and at 5, 15, 30, 60, 120, 240, 360, 480 and 1,440 min after intravenous administration. Three mice contributing plasma samples at each time point and each mouse sampled no more than twice. Blood was collected by submandibular sampling at non-terminal time points or by cardiac puncture for terminal sampling into lithium-heparin separator tubes. Samples were immediately placed on ice, and plasma was isolated by centrifugation and stored at −80 °C until analysis.

Plasma samples were prepared by protein precipitation with cold methanol following addition of tolbutamide as the internal standard. Following centrifugation, supernatants were evaporated, reconstituted, and analyzed by LC-MS. Calibration standards were prepared in blank mouse plasma over the validated concentration range used for sample analysis.

Chromatographic separation was performed using a Vanquish Binary liquid chromatography system (Thermo Fisher Scientific) equipped with an Uptisphere 120 Å ODB C18 column (3 µm, 2.1 × 30 mm) maintained at 40 °C. Samples (5 µl) were eluted at 0.3 ml/min using a 3.4-min gradient of 0.1% formic acid in water (mobile phase A) and acetonitrile (mobile phase B). ZIM was detected using a TSQ Quantis triple-quadrupole mass spectrometer (Thermo Fisher Scientific) operated in negative-ion selected-reaction monitoring mode, using transitions of m/z 424→58 and 424→265 for ZIM and 269→106 and 269→170 for the internal standard, tolbutamide. ZIM concentrations were determined from matrix-matched calibration curves based on analyte-to-internal-standard peak-area ratios. The lower limit of quantification was 19.5 nM.

Pharmacokinetic parameters were estimated by non-compartmental analysis of the composite mean plasma concentration–time profiles. The maximum observed plasma concentration (*C*_max_) and corresponding time (*T*_max_) were obtained directly from the observed group-mean concentration–time profiles. Variability at the (*C*_max_) time point was summarized using the sample standard deviation across the available replicate measurements. The area under the concentration-time curve from the start of dose administration to the last measurable time point, using the linear/log trapezoidal method (*AUC*_last_). For the oral group, the predose concentration was assumed to be zero, allowing inclusion of the interval from 0 to 15 minutes. For intravenous group, AUC was calculated from the first observed concentration at 5 min and therefore excludes the unobserved exposure between 0 and 5 min. The terminal elimination rate constant (*λ*_z_) was estimated by unweighted least-squares regression of the natural logarithm of the group-mean plasma concentration against time. The estimated oral bioavailability was calculated.

### ZIM Measurement in Tumor Mouse by HPLC/MS/MS

Methanol was added to samples at a ratio of 40µl per mg for tissue samples and 1/20 for serum samples (volume ratio, serum/methanol). The mixture was vortexed, homogenized (12 pulses at 10) and incubated on ice-bath for 30mins and the mixture was vortexed every 5 minutes during the incubation, followed by centrifugation at 10,000 rpm for 15 mins at 4 ℃. The supernatant was collected and the extraction procedure was repeated one more time. The combined supernatants were dried by speed-Vac and then re-dissolved in 100µl of methanol/acetonitrile (50/50, V/V) prior to HPLC/MSMS analysis.

An Agilent 1200 series HPLC system coupled to a 3200 QTRAP mass spectrometer (AB Sciex, Concord, ON, Canada) and using Analyst 1.4.2 software for data acquisition and analysis. Chromatographic separation was performed on a YMC Carotenoid column (10 cm × 2.0 mm ID; YMC America Inc.) under isocratic conditions with 90% methanol and 10% water at a flow rate of 200 µl/min. The injection volume was 5 µl.

Analytes were detected using an electrospray (ESI) source operated in negative ion mode. Quantification of ZIM was achieved using multiple reaction monitoring (MRM) with transition ion pairs 424>58 and 424>265. These transitions were selected based on product ion scans of the synthesized ZIM standard (molecular weight 425; **Supplementary Fig. S14**). Identification of ZIM in samples was confirmed by matching both MRM transitions and retention time to the in-house synthesized standard.

A seven-point calibration curve was generated using ZIM standards spiked with the internal standard at concentrations of 73.5, 147, 294, 1471, 2941, 7353, and 14706 nM. Calibration was based on the peak area ratio of ZIM to the internal standard plotted against ZIM concentration. Sample concentrations were calculated from this calibration curve.

### Off-target Profiling of ZIM by InVEST44

Off-target activity of ZIM was evaluated using the InVEST44 panel, comprising 44 pharmacologically relevant targets, including G protein–coupled receptors (GPCRs), ion channels, transporters, kinases, phosphodiesterases, cyclooxygenases, cholinesterases, monoamine oxidases, and nuclear receptors. Assays were conducted using target-appropriate binding or functional formats as defined by the platform provider, including radioligand binding, fluorescence polarization, electrophysiology, FLIPR-based assays, and cell-based reporter assays. ZIM was tested at a final concentration of 1 μM in duplicate. Reference compounds were included for each target and tested in full concentration–response format according to platform standards. Data are reported as percent activity relative to baseline control. For all assays except agonist mode, values approaching 100% indicate no target engagement, whereas lower values reflect increasing inhibition. In agonist model, 0% indicate no effect of ZIM, whereas higher values reflect increasing induction.

### Patients-Derived Organoid Generation

Prostate cancer tissues were collected with written informed consent obtained confidentially from all patients, according to the University of Alberta Hospitals Review Broad Committee approved protocol (HREBA.CC-14-0085). Tumor samples were immediately transferred to sterile culture medium and transported on ice to the laboratory for organoid generation within one hour of collection. Tumor tissues were first minced using sterile scalpel blades into 0.5-1 mm^3^, then digested by 5 mg/mL collagenase type II (Thermo Fisher Scientific, 17101015) and 0.5 mg/mL pronase (Sigma-Aldrich, 10165921001) in Advanced DMEM/F12 (Gibco, Thermo Fisher Scientific, 12634010) with 10 μM Y-27632 ROCK inhibitor (Cayman Chemical, 129830-38-2) and sterilely incubated at 37°C with 5% CO2 while gently stirred for 60-90 min. Dissociated cells were passed through a 40 µm cell strainer and washed with the Advanced DMEM/F12 medium. For organoid culture, a master mix at a ratio of 20,000 cells/40 µL growth factor in reduced Matrigel (Corning, 354230) was plated in the center of each well of a prewarmed 24-well plate, which was placed upside-down in a 37℃ tissue culture incubator and droplets allowed to solidify for 20 min. Matrigel droplets were overlayered with Advanced DMEM/F12 organoid growth media (OGM) containing 10 μM, SB202190 (Cayman Chemical, 152121-30-7), 1 μM prostaglandin E2 (Cayman Chemical, 363-24-6), 500 nM A83-01 (Cayman Chemical, 909910-43-6)), 10 ng/mL FGF-10 (PeproTech, Thermo Fisher Scientific, 100-26-25UG), 25 ng/mL EGF (PeproTech, Thermo Fisher Scientific, AF-100-15-500UG), 5 ng/mL FGF-2 (PeproTech, Thermo Fisher Scientific, 100-18B-50UG), 1 nM 5-alpha-dihydrotestosterone DHT (Sigma-Aldrich, D073), 1.25 mM N-acetyl-cysteine (Sigma-Aldrich, 616-91-1), 10 mM Nicotinamide (Cayman Chemical, 98-92-0), 1 × B-27 supplement (Gibco, Thermo Fisher Scientific, 17504044) and 20% conditioned medium of L-WRN cells, which are derived from stably transfecting L-Wnt3a cells with a R-spondin 3 and noggin co-expressing vector(59). Importantly, 10 µM Y-27632 ROCK inhibitor was added during the first week of growth and after passaging to promote growth. The medium was changed every 2-3 days.

### Electrophoretic mobility shift assay (EMSA)

ZNF281–DNA binding following ZIM treatment was assessed by electrophoretic mobility shift assay (EMSA). The IRDye 800 end-labeled ZNF281 oligonucleotides (sense: 5′-ATGGGGCGGGGTGGGGGG-3′; antisense: 5′-ATCCCCCCACCCCGCCCC-3′) were designed for this study and synthesized, infrared-labeled, and purified by Integrated DNA Technologies (IDT). Prior to use, complementary oligonucleotides were annealed by heating at 100 °C for 5 min in a heat block, followed by gradual cooling to room temperature. The binding reaction was performed by incubating 5 μg of lysates containing overexpressed ZNF281 protein with the infrared dye–labeled DNA probe using the Odyssey EMSA Kit (LI-COR Biosciences, 829-07910) for 20 minutes at room temperature, according to the manufacturer’s instructions. The reaction mixtures were resolved on 5% TBE polyacrylamide gels (Bio-Rad, 4565015) in 0.5× TBE buffer at 70 V for 2 hours under dark conditions.

### Statistical analysis

Use of parametric tests was decided after assessing the normal distribution of the values by normality tests. For parametric analysis, the unpaired t-test was used to compare two groups. One-way ANOVA and post hoc multiple-comparison correction was used to compare multiple groups. For non-parametric analysis, the Mann-Whitney U test was used for comparisons between two groups, and the Kruskal-Wallis test with Dunn’s multiple comparisons test correction was used for comparisons among several groups. Non-parametric Spearman’s rank correlation coefficient or the Pearson correlation coefficient was used for correlation analysis. The specific statistical test applied to each experiment is indicated in the respective figure legend. Statistical significance was defined by *P* < 0.05.

## Acknowledgments

The authors thank Dr. Lynne Postovit’s lab for providing the male NSG mice. We appreciate Dr. David Evans and Nicole Favis for supporting the usage of the IVIS machine. The Centre for Health Genomics and Informatics at the University of Calgary performed RNA and ChIP sequencing work. All the schematics were created using BioRender.com.

## Author contributions

G.H. generated the hypothesis, designed the study and performed all *in vivo* and mechanistic experiments, along with writing the manuscript. M.A.L.C. performed bioinformatic analysis of the sequencing data and some *in vivo* experiments. H.C. performed organoid culture and staining. R.C contributed to some in-vivo experiments. S.A.D, J.N, D. R. performed some mechanistic experiments. A. H. performed confocal microscope imaging. Y.Y.Z. performed mass spectrometry experiments and analysis. R.M. contributed to the pathologic analysis of the patients’ samples.

J.R.U. contributed to ZNF281 drug development. R.B.M., E.D.M., and J.L. coordinated mechanistic experiments and supervised students. A.T.D. developed, generated, and validated the ZNF281 inhibitor ZIM. A.K. and G.S. generated the hypothesis, designed the study, secured funding, coordinated experiments, supervised students, contributed to the ZNF281 drug development, and co-wrote the manuscript. All authors edited the manuscript.

## Data and materials availability

Raw sequencing data have been deposited in the NCBI Sequence Read Archive (SRA) under BioProject accession PRJNA1427223. Processed ChIP-seq data, including normalized bigWig signal tracks and MACS2 peak calls, have been deposited in the Gene Expression Omnibus (GEO) under accession GSE342271. The data will be released publicly upon publication. All code used for data preprocessing, statistical analyses, functional enrichment, pathway inference, and figure generation was written in R and relies exclusively on open-source software packages. Correspondence and requests for materials should be addressed to Adam Kinnaird and Gopinath Sutendra.

