## Supplementary Figures and Tables for "Pharmacologic Targeting of ZNF281 Suppresses Metastatic Prostate Cancer Beyond Androgen Receptor Dependence"

### Supplementary Figures & Tables

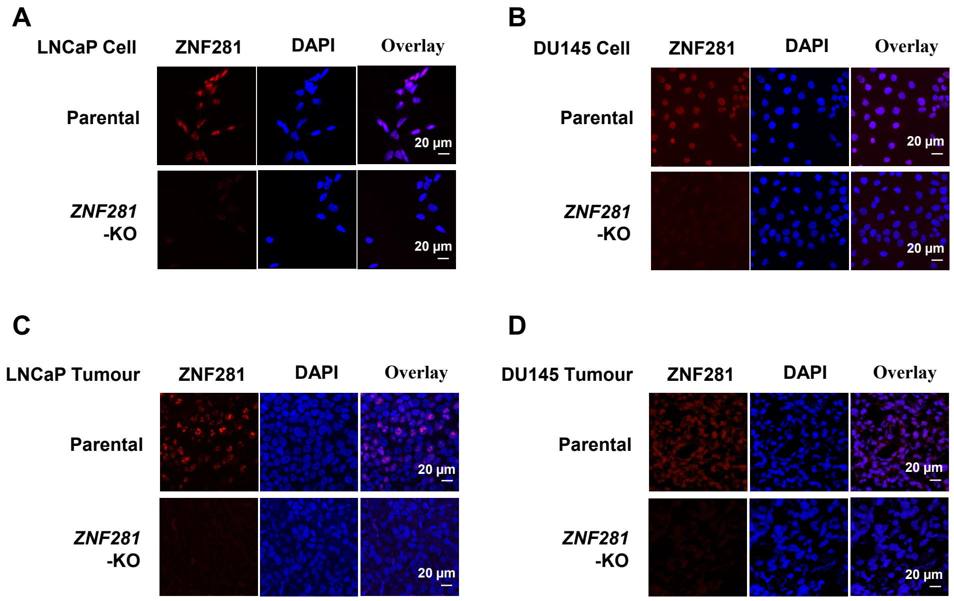


**Fig. S1. Validation of ZNF281 knockout in prostate cancer cell lines and orthotopic xenograft tumours by immunofluorescence staining. A, B,** Representative immunofluorescence images showing ZNF281 expression (red) in parental and *ZNF281*-KO LNCaP **(A)** and DU145 **(B)** cells. Nuclei were counterstained with DAPI (blue). Merged images are shown in the right panels. **C, D,** Representative immunofluorescence staining of ZNF281 (red) in orthotopic xenograft tumours derived from parental and *ZNF281*-KO LNCaP **(C)** and DU145 **(D)** cells. Nuclei were counterstained with DAPI (blue). Merged images are shown in the right panels. Scale bars, 20 μm.

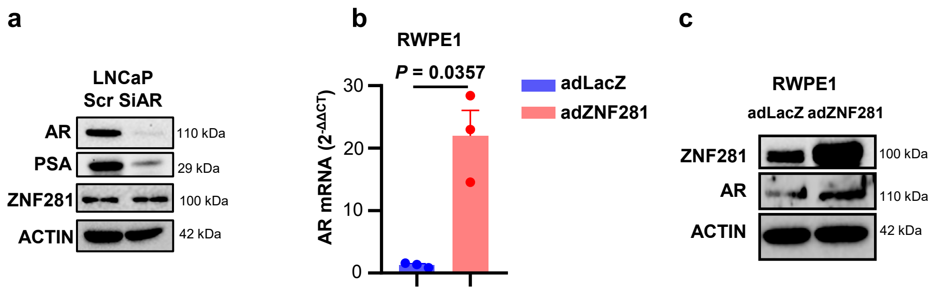


**Fig. S2. Regulatory relationship between AR and ZNF281. A,** Western blot analysis showing that AR knockdown robustly reduced the expression of its known target gene PSA, while having no inhibitory effect on ZNF281 protein expression. **B,** RT-qPCR analysis showing that overexpression of ZNF281 increased AR mRNA expression in the immortalized normal prostate epithelial cell line RWPE1. **C,** Western blot analysis showing that overexpression of ZNF281 increased AR protein expression in the immortalized normal prostate epithelial cell line RWPE1.

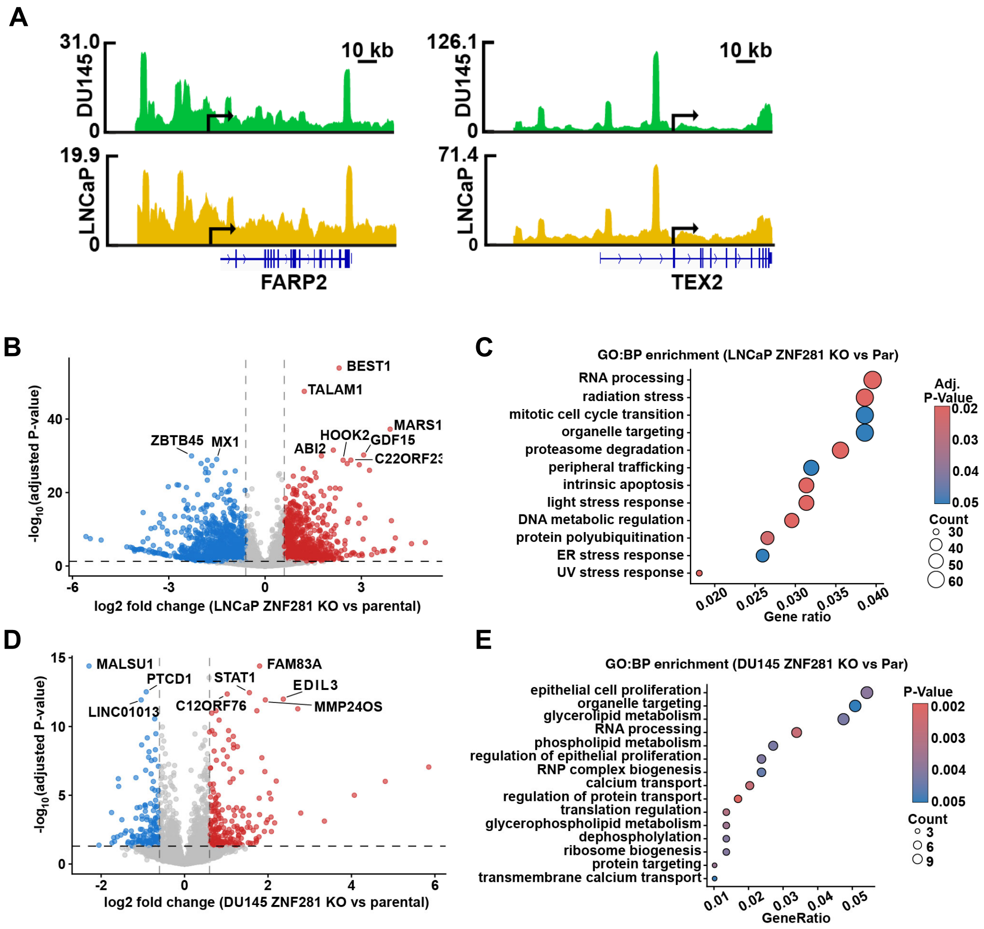


**Fig. S3. Functional consequences of ZNF281 loss in LNCaP and DU145 cells. A,** Genome browser tracks showing ZNF281 ChIP-seq signal in DU145 (green) and LNCaP (yellow) cells at representative shared DEGs (FARP2, and TEX2). Tracks illustrate differential ZNF281 promoter occupancy across cell lines. Gene models and transcriptional orientation are shown below, with scale bars indicating genomic distance. **B,** Volcano plot showing differential gene expression in LNCaP cells comparing ZNF281 knockout (KO) to parental cells. Genes significantly upregulated or downregulated are highlighted, based on adjusted P value and $log_2$ fold-change thresholds. Selected genes of interest are annotated. **C,** Gene Ontology (GO) biological process enrichment analysis of differentially expressed genes in LNCaP ZNF281 KO versus parental cells. Enriched pathways are shown as a dot plot, with dot size representing the number of genes contributing to each pathway and color indicating statistical significance (*P* value). **D,** Volcano plot showing differential gene expression in DU145 cells comparing ZNF281 knockout (KO) to parental cells. Genes significantly upregulated or downregulated are highlighted, based on adjusted P value and $log_2$ fold-change thresholds. Selected genes of interest are annotated. **E,** Gene Ontology (GO) biological process enrichment analysis of differentially expressed genes in DU145 ZNF281 KO versus parental cells. Enriched pathways are shown as a dot plot, with dot size representing the number of genes contributing to each pathway and color indicating statistical significance (*P* value).

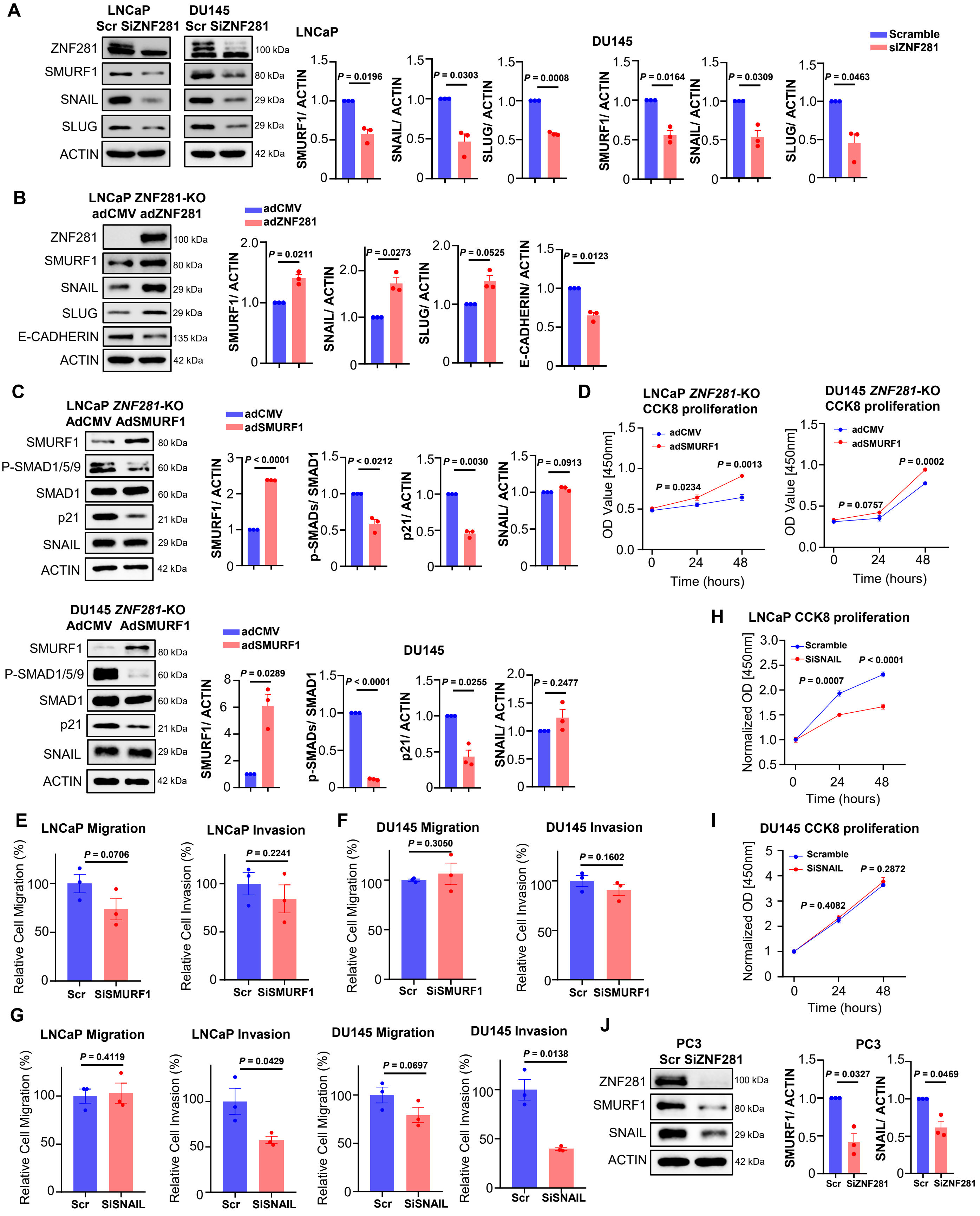


**Fig. S4. AR-independent functions of ZNF281 in prostate cancer progression involve SMURF1-mediated growth and SNAIL-mediated EMT. A,** Representative western blots and quantification showing the effects of siRNA-mediated ZNF281 knockdown on SMURF1, SNAIL and SLUG in LNCaP and DU145 cells. **B,** Representative western blots and quantification showing the effects of adenoviral ZNF281 re-expression on SMURF1 and EMT-related proteins including SNAIL, SLUG, and e-cadherin in ZNF281-KO LNCaP cells. **C,** Representative western blots and quantification showing the effects of adenoviral SMURF1 re-expression on the SMURF1–p-SMAD1/5/9–p21 axis and SNAIL in ZNF281-KO LNCaP and DU145 cells. **D,** CCK8 assays showing the effects of SMURF1 re-expression on the proliferation of ZNF281-KO LNCaP and DU145 cells. **E, F,** Transwell migration and invasion assays following siRNA-mediated SMURF1 knockdown in LNCaP (E) and DU145 (F) cells. **G,** Transwell migration and invasion assays following siRNA-mediated SNAIL knockdown in LNCaP and DU145 cells. **H, I,** CCK-8 assays showing the effects of siRNA-mediated SNAIL knockdown on the proliferation of LNCaP (H) and DU145 (I) cells. **J,** Representative western blots and quantification showing the effects of siRNA-mediated ZNF281 knockdown on SMURF1 and SNAIL protein abundance in PC3 cells. Data are presented as the mean ± SEM from three independent biological replicates. Statistical significance was determined using t test with Welch's correction.

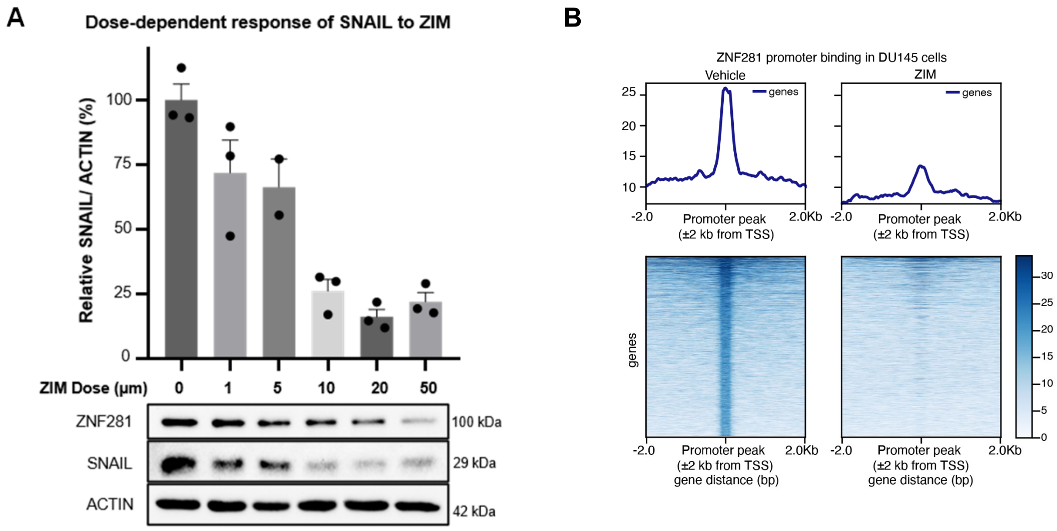


**Fig. S5. Generation of the orally bioavailable ZNF281 Interfering Molecule (Oral ZIM). A,** Western blot analysis and quantification showing the dose-dependent effects of oral version ZIM treatment on ZNF281-regulated protein SNAIL in DU145 cells. Protein expression was normalized to ACTIN. 10 μM ZIM achieved markedly inhibition of SNAIL expression, was used in subsequent in vitro mechanistic experiments to achieve robust target-pathway inhibition. **B,** Metaplot (top) and heatmap (bottom) of ZNF281 ChIP-seq signal at promoter regions (±2 kb from TSS) in DU145 cells under vehicle or ZIM treatment, demonstrating ZIM-induced loss of ZNF281 promoter occupancy in a second cellular context. Data are presented as the mean ± SEM from three independent biological replicates. Statistical significance was determined using t test with Welch's correction.

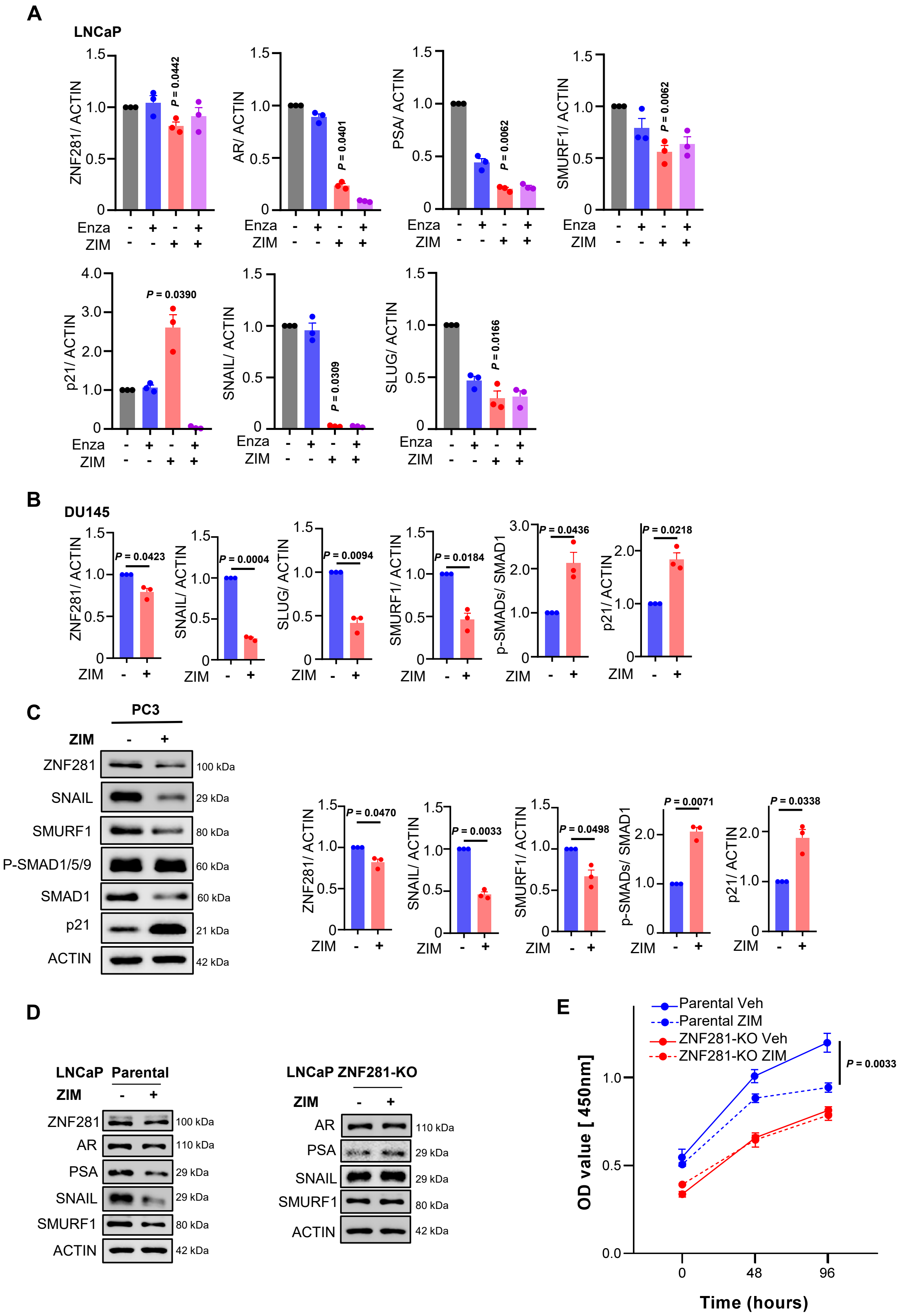


**Fig. S6. Western blot and quantification of oral version ZIM treatment in LNCaP, DU145, and PC3 cells. (A)** Western blot quantification showing the effects of ZIM (10 µM) and enzalutamide (10 µM) in LNCaP cells. **(B)** Western blot quantification showing the effects of ZIM (10 µM) in DU145 cells. **(C)** Representative western blot and quantification of PC3 cells treated with ZIM (10 µM) for 48 hours. **(D)** Western blot analysis of the effects of ZIM treatment (1 µM) in LNCaP parental and ZNF281-KO cells. **(E)** In vitro CCK8 proliferation assay showing the antiproliferation effect of ZIM treatment (1 µM) in LNCaP parental and ZNF281-KO cells. Graphs show the mean ± SEM from three independent biological replicates. Statistical significance was determined using Kruskal-Wallis test or t test with Welch's correction.

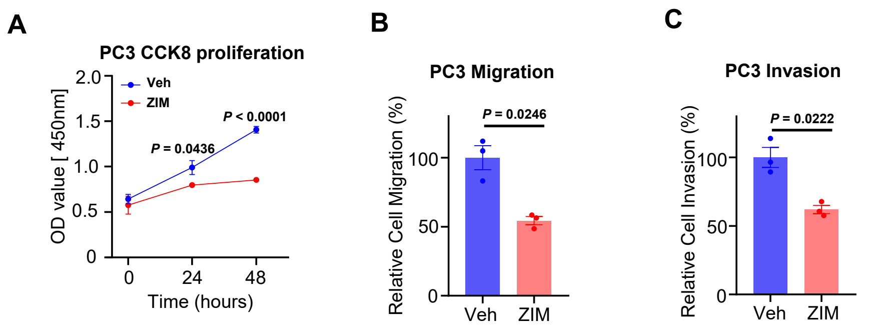


**Fig. S7. ZIM effects on PC3 cells. (A)** CCK-8 assays showing the effect of ZIM treatment on PC3 cell proliferation over 48 h. **b, c,** Transwell assays showing the effects of ZIM on PC3 cell migration **(B)** and invasion **(C)**. Graphs show the mean ± SEM from three independent biological replicates. Statistical significance was determined using t test with Welch's correction.

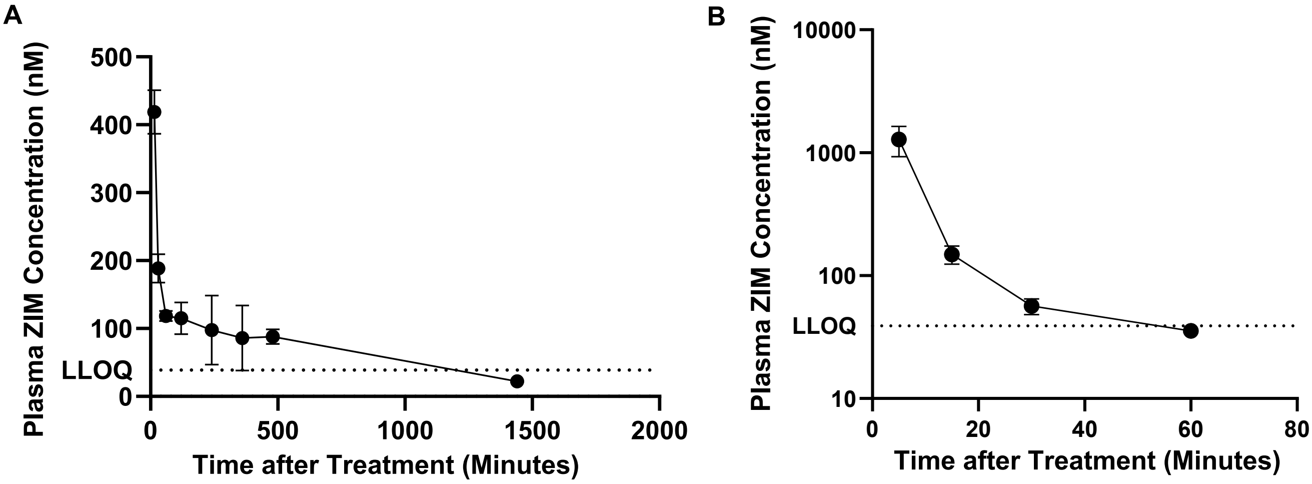


**Fig. S8. Pharmacokinetic study of ZIM following oral and intravenous administration.** Plasma concentrations of ZIM following a single oral dose of 250 mg/ kg or intravenous dose of 2.5 mg/ kg in mice. Blood samples were collected at 15, 30, 60, 120, 240, 360, 480, 1440 min after oral administration **(A)** and at 5, 15, 30, 60, 120, 240, 360, 480 min after intravenous administration **(B)**. Data are presented as the mean ± SD from three independent biological replicates. Concentrations below the lower limit of quantification are indicated as LLOQ.

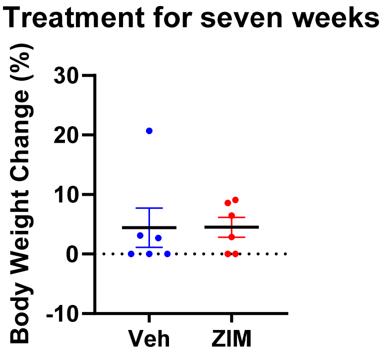


**Fig. S9. Body weight changes after ZIM treatment.** Percentage changes in body weight relative to the pretreatment baseline were monitored in healthy, non-tumour-bearing mice receiving vehicle or ZIM (250 mg/kg once daily by oral gavage for seven weeks). Data are presented as mean ± SEM.

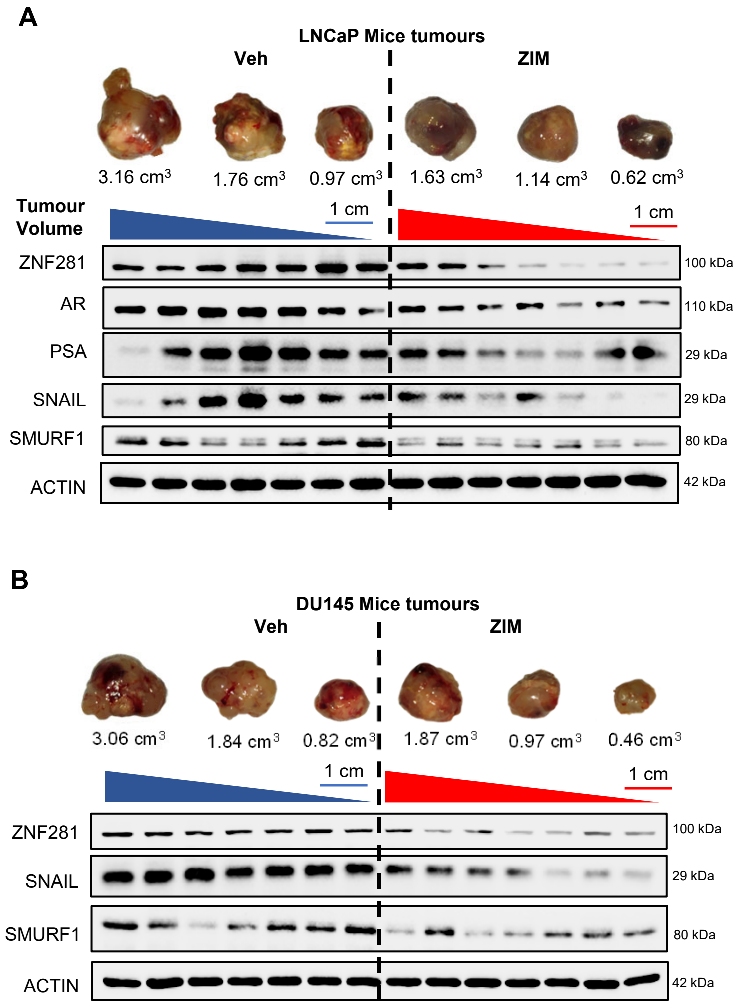


**Fig. S10. Effects of ZIM on ZNF281-associated signaling in orthotopic xenograft tumours.** Western blot analysis of ZNF281 and the indicated downstream proteins in individual vehicle and ZIM treated LNCaP **(A)** and DU145 **(B)**. The largest, median, and smallest tumours in each group were shown. Tumours in each group were arranged from largest to smallest.

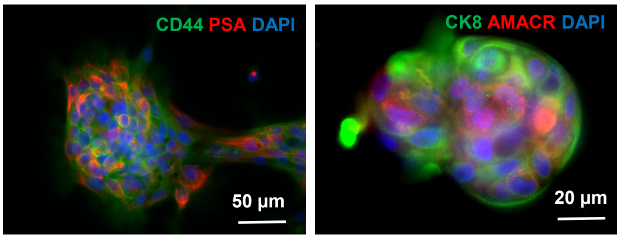


**Fig. S11. Representative immunofluorescence images of patient-derived organoids.**

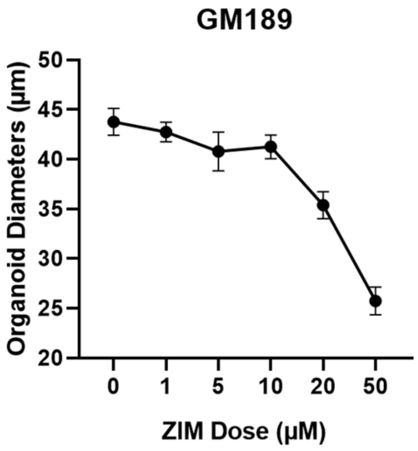


**Fig. S12. ZIM reduced organoids size in a dose-dependent manner.** Diameter of GM189 patient-derived organoids following treatment with the indicated concentrations of ZIM for 7 days. Data are presented as mean ± SEM.

A

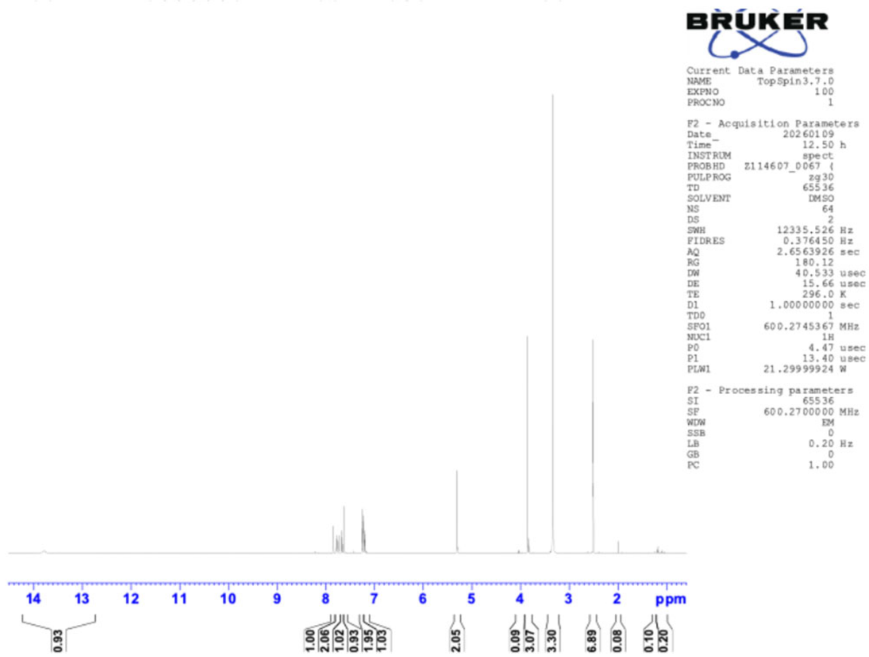


B

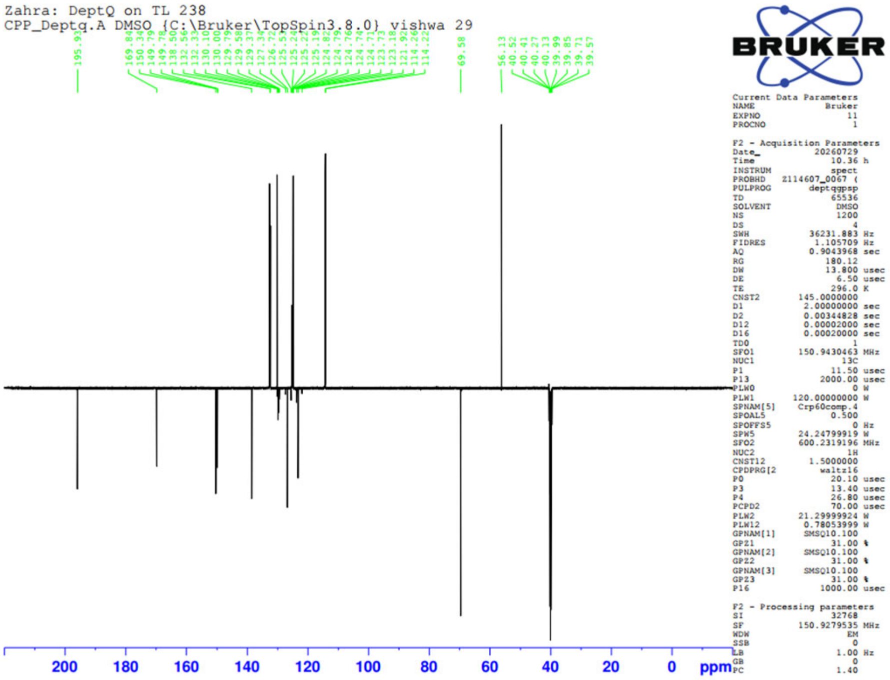


C

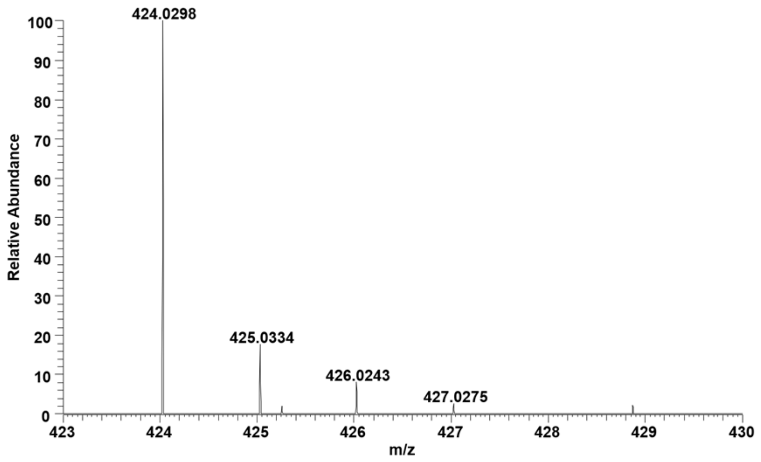


**Fig. S13. $^{1}$H NMR spectra (A), $^{13}$C NMR spectra (B), and HRMS (C) of ZIM.**

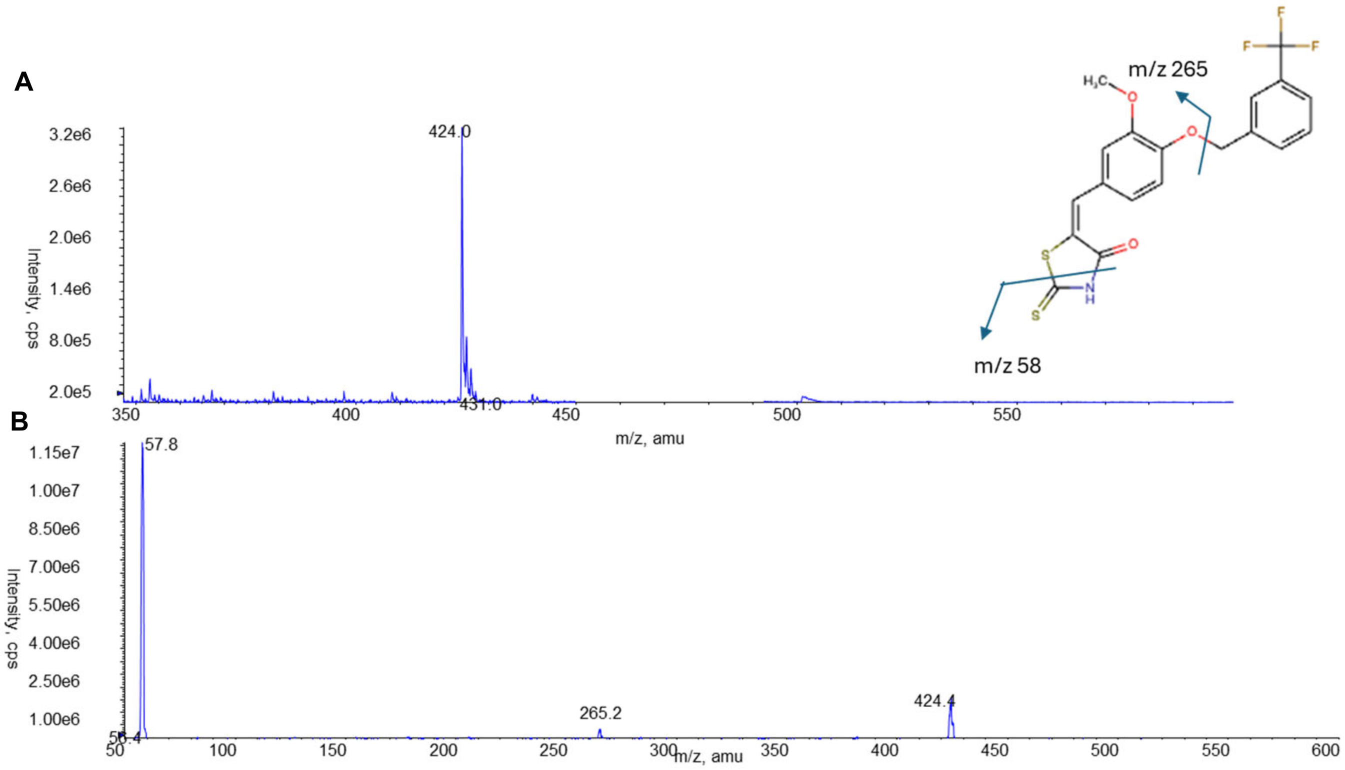


**Fig. S14. HPLC-MS analysis of ZIM. (A)** Full scan mass spectrum of the compound, showing a prominent molecular ion peak at *m/z* 424.0. **(B)** MS/MS fragmentation spectrum of the ion at *m/z* 424.0 ion, showing major fragment ions at *m/z* 265.2 and *m/z* 58.0. Fragment assignments correspond to cleavage of specific structure, as illustrated in the figure.

**Clinical Characteristics of the Fifteen Patients**

| Patient | Age | PSA (ng/ml) | Gleason Grade Group | TNM |
|---|---|---|---|---|
| U17-7679 | 62 | 7.1 | 3 | T3bN1M0 |
| U16-19109 | 69 | 21.4 | 2 | T3bN1M0 |
| RH18-25230 | 58 | 16.4 | 2 | T3bN1M0 |
| U19-20443 | 66 | 6.8 | 3 | T3aN1M0 |
| RAS20-004052 | 55 | 5.5 | 3 | T3bN1M0 |
| U18-24783 | 57 | 3.3 | 5 | T3aN1M0 |
| RH17-25850 | 57 | 25.2 | 2 | T3aN1M0 |
| RH17-25234 | 70 | 7.4 | 5 | T3bN1M0 |
| RH18-3925 | 64 | 9 | 2 | T3aN1M0 |
| U18-23320 | 60 | 10 | 2 | T3bN1M0 |
| RH19-8985 | 54 | 14.7 | 3 | T3aN1M0 |
| RH18-165620 | 68 | 8.5 | 2 | T3bN1M0 |
| RH19-7813 | 68 | 14.9 | 3 | T3aN1M0 |
| RH19-9364 | 60 | 6 | 3 | T3aN1M0 |
| RE-U17-16206 | 57 | 10.2 | 3 | T3bN1M0 |

*AJCC 8th Edition

**Table S1. Clinical characteristics of the fifteen patients for IF staining.**

| Target Family | Target | % Activity ZIM Data 1 | ZIM Data 2 | IC50 (M) Reference Compound (EC50 for agonist) | Reference Compound | Radioligand (RL) (all [3H]-ligands) | RL Conc. (nM) |
|---|---|---|---|---|---|---|---|
| G-protein Coupled Receptors (GPCRs) | 5-HT1A | 104 | 102 | 9.36E-09 | 8-OH DPAT | 8-OH DPAT | 10 |
| | 5-HT1B | 100 | 99 | 7.71E-09 | GR125743 | GR125743 | 10 |
| | 5-HT2A | 112 | 116 | 5.12E-10 | Ketanserin | LSD | 2 |
| | 5-HT2B | 86 | 87 | 3.47E-08 | Serotonin | LSD | 2 |
| | Adenosine A2A | 48 | 47 | 8.15E-08 | CGS212680 | CGS21680 | 15 |
| | Adrenergic a1A | 96 | 96 | 3.12E-09 | Prazosin | Prazosin | 10 |
| | Adrenergic a2A | 97 | 99 | 8.62E-09 | RX 821002 | RX821002 | 5 |
| | Adrenergic b1 | 101 | 100 | 3.03E-09 | Alprenolol | Alprenolol | 1 |
| | Adrenergic b2 | 104 | 103 | 6.12E-09 | Alprenolol | Alprenolol | 1 |
| | Cannabinoid CB1 | 106 | 106 | 4.61E-09 | CP55,940 | CP55,940 | 1 |
| | Cannabinoid CB2 | 95 | 93 | 2.96E-09 | WIN55212-2 | CP55,940 | 2 |
| | CCK1 | 110 | 118 | 1.11E-07 | Lorglumide | CCK | 5 |
| | Dopamine D1 | 98 | 99 | 2.55E-09 | SCH 23390 | SCH 23390 | 0.3 |
| | Dopamine D2S | 102 | 101 | 6.65E-10 | Spiperone | Methylspiperone | 0.2 |
| | Endothelin ETA (Agonist) | 0 | 0 | 1.20E-07 | Endothelin | | |
| | Endothelin ETA (Antagonist) | 88 | 93 | 2.86E-08 | Sitaxentan | | |
| | Histamine H1 | 100 | 105 | 6.19E-09 | Mepyramine | Pyrilamine | 5 |
| | Histamine H2 | 87 | 95 | 1.65E-08 | Ranitidine | Famotidine | 20 |
| | Muscarinic M1 | 101 | 103 | 7.80E-09 | Pirenzepine | Pirenzipine | 5 |
| | Muscarinic M2 | 115 | 114 | 1.27E-08 | AF-DX 384 | AF-DX 384 | 5 |
| | Muscarinic M3 | 101 | 102 | 4.43E-09 | 4-DAMP | NMS | 0.2 |
| | δ opioid | 95 | 95 | 1.51E-08 | DADLE | DADLE | 10 |
| | κ opioid | 97 | 95 | 1.40E-08 | U-69,593 | U69,593 | 15 |
| | μ opioid | 98 | 97 | 2.07E-08 | DAMGO | DAMGO | 15 |
| | Vasopressin V1A (Agonist) | -1 | -1 | 4.64E-12 | Vasopressin | | |
| | Vasopressin V1A (Antagonist) | 96 | 91 | 1.19E-08 | SR-49059 | | |
| Transporters | DAT | 104 | 108 | 1.45E-07 | AHN 1-055 | WIN35428 | 30 |
| | NET | 134 | 130 | 8.18E-09 | Nisoxetine | Nisoxetine | 4 |
| | SERT | 78 | 87 | 6.26E-08 | Imipramine | Imipramine | 20 |
| Ion Channels | 5HT3 | 106 | 101 | 1.35E-08 | Quipazine | Ondansetron | 20 |
| | GABA-A Central BZD | 86 | 89 | 3.71E-09 | Flumazenil | Ro151788 | 10 |
| | CaV1.2 | 95 | 96 | 8.85E-08 | Nifedipine | | |
| | hERG | 91 | 96 | 2.35E-08 | E-4031 | | |
| | NaV1.5 | 93 | 97 | 8.92E-07 | TTX | | |
| | Nicotinic AChR a4b2 (Agonist) | 1 | -2 | 1.18E-08 | Epibatidine | | |
| | Nicotinic AChR a4b2 (Antagonist) | 109 | 103 | 5.33E-07 | Mecamylamine | | |
| | NMDA | 100 | 100 | 1.64E-08 | MK-801 | MK-801 | 10 |
| Kinase | Lck | 78 | 74 | 9.85E-10 | Staurosporine | | |
| Phosphodiesterases | PDE3A | 107 | 107 | 1.96E-08 | Cilostamide | | |
| | PDE4D2 | 101 | 100 | 1.11E-07 | Rolipram | | |
| Cyclooxygenases | COX-1 | 104 | 98 | 5.23E-08 | SC-560 | | |
| | COX-2 | 104 | 102 | 1.00E-07 | DuP-697 | | |
| Cholinesterases | Acetylcholinesterase | 108 | 110 | 6.96E-08 | Physostigmine | | |
| Monoamine Oxidases | MAO-A | 122 | 123 | 1.64E-07 | Tranylcypromine | | |
| | MAO-B | 51 | 52 | 6.09E-08 | Tranylcypromine | | |
| Nuclear Receptors | Androgen Receptor (Agonist) | 0 | 0 | 4.38E-11 | 5aDH-11-kT | | |
| | Androgen Receptor (Antagonist) | 62 | 64 | 5.88E-08 | Apalutamide | | |
| | Glucocorticoid Receptor | 104 | 102 | 1.61E-08 | Dexamethasone | | |

**Table S2. Off-target profiling of ZIM across a panel of pharmacologically relevant targets (InVEST44 platform).** ZIM was evaluated against a panel of 44 targets encompassing G protein–coupled receptors (GPCRs), ion channels, biogenic amine transporters, nuclear receptors, phosphodiesterases, kinases, and proteases using a combination of radioligand binding, fluorescence polarization, electrophysiology/FLIPR, and reporter-based assays. The compound was tested at a final concentration of 1 μM in duplicate. Data are presented as percent activity relative to baseline. For all assays except agonist mode, values approaching 100% indicate no target engagement, whereas lower values reflect increasing inhibition. In agonist model, 0% indicate no effect of test compound, whereas higher values reflect increasing induction. Across the panel, ZIM exhibited minimal off-target activity, with the majority of targets showing activity within 75–125% or around 0% (agonist model) of baseline, consistent with negligible modulation.

| PK parameter | Oral, 250 mg/kg | IV, 2.5 mg/kg |
|---|---|---|
| $C_{max}$ (nM) | 419.1 ± 56.0 | 1283.7 ± 613.8 |
| $T_{max}$ (min) | 15 | 5 |
| $C_{last}$ (nM) | 22.3 | 35.5 |
| $T_{last}$ (min) | 1440 | 60 |
| $t_{1/2}$ (min) | 554.2 | 23.5 |
| $AUC_{last}$ (nM·min) | 106515 | 10084 |
| $AUC_{0\text{-}\infty}$ (nM·min)[a] | 124316 | 11285 |
| *AUC* extrapolated (%) | 14.3 | 10.6 |
| λz ($min^{-1}$) | 0.00125 | 0.0295 |
| Estimated Oral bioavailability, *F* (%) | 10.6 | — |

[a] For the intravenous group, extrapolation to time zero is limited by the absence of sampling prior to 5 minutes post-dose. Therefore, $AUC_{0\text{-}\infty}$ was interpreted with this limitation.

**Table S3. Pharmacokinetic parameters of ZIM following oral and intravenous administration.** Plasma concentrations of ZIM following a single oral dose of 250 mg/ kg or intravenous dose of 2.5 mg/ kg in mice. Blood samples were collected at 15, 30, 60, 120, 240, 360, 480, 1440 min after oral administration and at 5, 15, 30, 60, 120, 240, 360, 480 min after intravenous administration. The maximum observed plasma concentration ($C_{max}$) and the corresponding sampling time ($T_{max}$) were obtained directly from the observed group-mean concentration–time profiles. Variability at the $C_{max}$ time point was summarized using the sample standard deviation across the available replicate measurements. The area under the concentration-time curve from the start of dose administration to the last measurable time point, using the linear/log trapezoidal method ($AUC_{last}$). For the oral group, the predose concentration was assumed to be zero, allowing inclusion of the interval from 0 to 15 minutes. For intravenous group, AUC was calculated from the first observed concentration at 5 min and therefore excludes the unobserved exposure between 0 and 5 min. The terminal elimination-rate constant (λz) was estimated by unweighted least-squares regression of the natural logarithm of the group-mean plasma concentration against time. Data are presented as mean ± SD.

**ZIM safety test in mice.**

| | Control (n = 6) | ZIM (n = 6) | *P*-value |
|---|---|---|---|
| **RBC (M/μL)** | 6.73 ± 0.07 | 6.72 ± 0.35 | 0.26 |
| **WBC (K/μL)** | 0.6 ± 0.08 | 0.83 ± 0.21 | 0.43 |
| **Platelet Count (K/μL)** | 643.5 ± 18.21 | 641.4 ± 40.34 | 1.00 |
| **HGB (g/dL)** | 10.47 ± 0.13 | 10.82 ± 0.16 | 0.15 |
| **HCT (%)** | 32.50 ± 0.37 | 32.92 ± 1.17 | 0.41 |
| **MCV (fL)** | 48.50 ± 0.22 | 49.50 ± 1.12 | 1.00 |
| **MCH (pg)** | 15.53 ± 0.06 | 16.37 ± 0.93 | 0.98 |
| **MCHC (g/dL)** | 32.22 ± 0.09 | 33.05 ± 1.04 | 0.85 |
| **ALB (g/L)** | 25.60 ± 0.66 | 28.82 ± 0.83 | 0.03 |
| **ALT (U/L)** | 26.90 ± 3.37 | 29.12 ± 5.70 | 1.00 |
| **AST (U/L)** | 89.22 ± 9.02 | 98.76 ± 15.23 | 0.84 |
| **BUN (mmol/L)** | 7.98 ± 0.51 | 8.93 ± 0.65 | 0.23 |

Mean ± SEM

**Table S4. ZIM safety test in healthy, non-tumour-bearing mice.**

**ZIM levels in mice.**

| Serum (μM) | Tumour (ng/mg) |
|---|---|
| 0.90 ± 0.25 | 7.04 ± 1.86 |

Mean ± SEM

**Table S5. ZIM levels measured by HPLC-MS/MS analysis in tumor mice.**

| Patient | Age | PSA (ng/ml) | Gleason Grade Group | TNM |
|---|---|---|---|---|
| GM114 | 48 | 35.8 | 5 | T3bN0M0 |
| GM153 | 75 | 47.9 | 4 | T4N1M0 |
| GM161 | 80 | 54.4 | 3 | T2bNxM1b |
| GM189 | 75 | 92.9 | 4 | T3N1M0 |
| GMRP2 | 57 | 3.5 | 2 | T3aN0M0 |
| GMRP3 | 47 | 9.3 | 2 | T2N0M0 |
| GM515 | 59 | 6.4 | 2 | T2N0M0 |
| GM520 | 47 | 2.9 | 2 | T2N0M0 |
| GM556 | 67 | 6.2 | 2 | T3N0M0 |
| GM557 | 63 | 12.8 | 4 | T1N0M0 |

*AJCC 8th Edition

**Table S6. Patient clinical profiles for organoid establishment.**
